# Hippocampal CA1 represents action and reward events instantly compared to the superficial and deep layers of the lateral entorhinal cortex

**DOI:** 10.1101/2022.03.31.485431

**Authors:** Shogo Soma, Shinya Ohara, Satoshi Nonomura, Junichi Yoshida, Naofumi Suematsu, Eva Pastalkova, Yutaka Sakai, Ken-Ichiro Tsutsui, Yoshikazu Isomura

## Abstract

The entorhinal cortex (EC) is the main interface between the hippocampus and the neocortex. The EC plays a critical role in learning and memory. We investigated the neuronal representation of behavioral events during operant learning in the hippocampal-entorhinal circuit of head-fixed rats. Both CA1 and lateral entorhinal cortex (LEC) neurons develop task-related activities after learning. Among diverse task-related activities, we compared the transient peak activities that represent action and reward and found a distinct difference in the timing of behavioral event representation between CA1 and LEC. CA1 represents action and reward events in close to real-time, whereas both the superficial and deep layers of the LEC showed delayed representation of those events. Our results suggest that subpopulations exist within which CA1 and LEC neurons process the information in a different order from the anatomically defined hippocampal-entorhinal circuit.

## Introduction

The entorhinal cortex (EC) is the major interface between the hippocampus and the neocortex. Together with the hippocampus, the EC plays a crucial role in processing information from ongoing events i.e., the *when-where-what* information necessary for episodic memory. Previous anatomical studies have investigated the hippocampal-entorhinal circuit in detail and shown clear segregation of the hippocampal-input and output circuits within the EC layers in both rodents and primates (Witter et al., 2017; Ohara et al., 2018, 2021a, 2021b). EC neurons in the superficial layers, layers II and III, constitute the hippocampal-input circuits by projecting to the dentate gyrus (DG)/CA3 and CA1, respectively. This information is processed through the hippocampal trisynaptic circuit (EC layer II → DG → CA3 → CA1) and the direct EC (layer III)-hippocampal input is integrated within CA1 neurons. CA1 neurons, in turn, send this information back to the deep layers of the EC, principally layer V, via the hippocampal-output circuit. This “entorhinal-hippocampal-entorhinal pathway” is considered the main circuit that supports information processing between the hippocampus and the neocortex.

The EC is differentiated anatomically and functionally into the medial and lateral EC (MEC and LEC, respectively). The two subdivisions receive inputs from different sensory modalities, such as light and smell, from the neocortex and integrate them into multimodal information (Kerr et al., 2007; Nilssen et al., 2019; Doan et al., 2019). The hippocampus processes this information according to entorhinal integration to realize higher order brain functions e.g., learning, memory, recognition, and navigation. For example, the MEC processes spatial information in a comprehensive manner (grid cells: Fyhn et al., 2004; Hafting et al., 2005; Deshmukh et al., 2010; border cells: Savelli et al., 2008; Solstad et al., 2008; head direction cells: Sargolini et al., 2006; and object vector cells: Høydal et al., 2019). Such spatial information is converted into a specific spatial location encoded by each neuron as a place cell in the hippocampus, eventually representing the space (*where*) of the external world (Zhang et al, 2013; Moser et al., 2015; Sugar and Moser, 2019).

In contrast to the MEC, spatially modulated cells are essentially absent from the LEC. Instead of pure spatial information, the LEC processes other information, such as that concerning odors and objects, and is involved in diverse memory functions, including object information (Deshmukh and Knierim, 2011; Tsao et al., 2013; Wang et al., 2018), sensory/context association learning (Lu et al., 2013; Igarashi et al., 2014; Keene et al., 2016; Li et al., 2016; Pilkiw et al., 2017; Lee et al., 2021), episodic-like memory (Chao et al., 2016; Vandrey et al., 2020), and trace conditioning (Morrissey et al., 2012; Tanninen et al., 2013, 2015).

One of the LEC’s key functions is to represent temporal information in a comprehensive manner. LEC neurons encode elapsed time on a second-to-minute timescale by demonstrating ramping activities related to memory (Tsao et al., 2018). This temporal information as well as information from the MEC (Robinson et al., 2017) can be used to form the more specific time representing activity (*when*) in the hippocampus (time cell, Pastalkova et al., 2008; MacDonald et al., 2011). These types of time representations have also been observed in the human LEC and hippocampus (Montchal et al., 2019; Umbach et al., 2020).

Thus, the hippocampus is an adequate region that processes the space and time of an episodic memory by focusing on specific information in a spatiotemporal framework provided by the EC (Rueckemann and Buffalo, 2017; Buzsáki and Tingley, 2018; Sugar and Moser, 2019). Based on this accumulating evidence, the EC has been proposed to represent the spatial, sensory, context, and time information of the universal environment and the hippocampus apparently extracts specific aspects of this information from the entorhinal-hippocampal-entorhinal pathway. Thus, the episodic memory encoded in the hippocampus is thought to be information integrated from various kinds of information sources, i.e., *what* information representing environmental events is associated with *where* and *when* information encoded by place and time cells, respectively. In that case, how and when do the EC and hippocampus represent the event as it is experienced in real-time? Additionally, how differently do superficial and deep EC layers process event information in the entorhinal-hippocampal-entorhinal pathway?

In this study, we investigated the timing of neural representation of two distinct behavioral events related to learning in the entorhinal-hippocampal-entorhinal pathway, CA1 and superficial and deep LEC layers. To do so, we adopted the simplest operant task in which outcome (reward) was presented after rats voluntarily moved their forelimb (action) without external instructive cues (Soma et al., 2017). Episodic experiences can be effective even for such elementary learning through trial and error repetition. By using rats with their heads fixed instead of rats that can move freely, we could measure the timing of events (action/outcome) precisely while monitoring neural activities (Soma et al. 2017, 2019; Rios et al., 2019). We recorded CA1 and LEC neurons extracellularly from rats both prior to and following training in this task. We used these recordings to examine the relationship between behavioral events and spike activities on a subsecond scale.

## Results

### Self-paced, spontaneous left or right pedal pressing task

To determine the timing of neural representation for two distinct behavioral events related to learning in the entorhinal-hippocampal-entorhinal pathway, we adopted the simplest operant task: a self-paced, spontaneous left or right pedal pressing task (Figure 1). The rats used in our study had to manipulate left and right pedals with the corresponding forelimb in a head-fixed condition, which enabled us to monitor the accurate timing of events (action/outcome). The rats started each trial spontaneously by pushing both pedals down with the left and right forelimbs and holding them for a constant period in a self-paced manner. After completing the holding period, the rats had to choose either the left or right pedal without any instructional cue indicating which pedal would provide the reward, and release the correct pedal (action) to obtain saccharin water as a reward (outcome). The task consisted of two blocks, right pedal rewarded and left pedal-rewarded blocks, and the reward pedal was changed in a block-by-block manner with no instruction (R, R, R, R… L, L, L, L…; Figure 1A and B).

**Figure 1.**
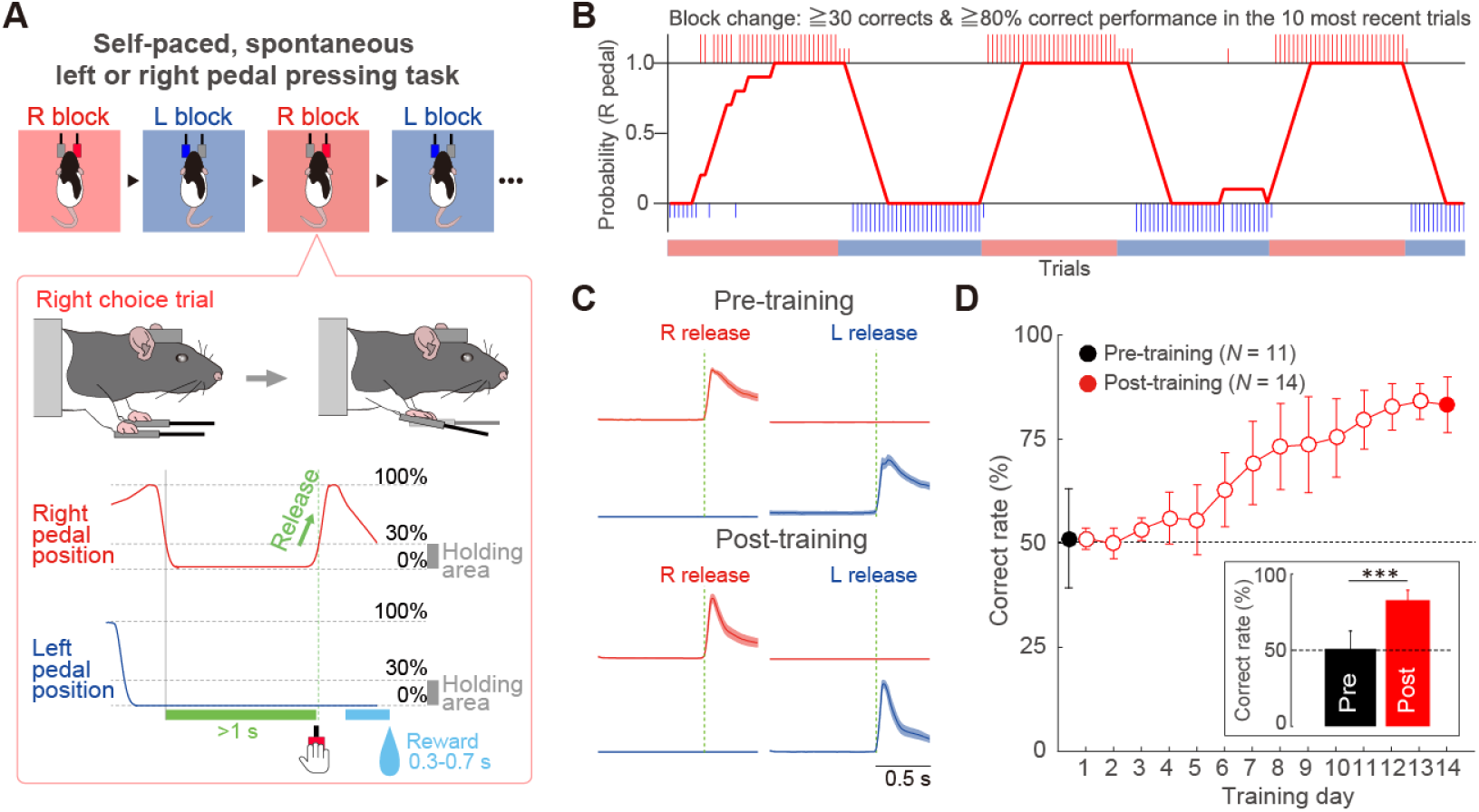
Behavioral task performance. (A) Schematic diagram of the self-paced, spontaneous left or right pedal pressing task. Head-fixed rat pushed down both pedals for a short period (≥1 s) to start each trial, and subsequently released either pedal (e.g., right release) voluntarily (without instructive cue) to acquire a reward. This task consisted of right-rewarded (R) and left-rewarded (L) blocks, which were alternated after meeting the criteria (see Methods). (B) A typical example of task performance. Rat chose the correct pedal based on the reward. Large and small colored vertical bars (red represents right choice; blue represents left choice) indicate correct and incorrect trials, respectively. We averaged the number of right correct choices obtained from the past 10 trials to calculate the proportion of correct choices. (C) Right-left pedal trajectories obtained from pre-(1^st^ day, top) and post-trained (14^th^ day, bottom) rats. (D) Learning curve over 14 training days. Inset, averaged proportion of correct choices (mean ± SD) in the 1^st^ and 14^th^ days. Black and red colors represent the pre- and post-training groups, respectively. *** *p* < 0.001, Mann–Whitney test.

Figure 1C show the pedal traces on the left- or right-releasing trials obtained from pre- and post-training groups. Both groups could manipulate the individual pedals spontaneously without left-right bias (see also Figure S1A). Thus, rats could manipulate the pedal i.e., they could spontaneously express the motor response (unilateral pedal release), before they learned the task rule. In contrast, other measures such as holding stability and release time (time from onset to end of release) were developed over the course of training. Rats in the post-training group quickly released the pedal and returned their forelimb to the pedal after stable pedal holding (Figure S1B), indicating that the rats learned the precise motor response (skilled pedal manipulation) over the course of training.

The rats typically learned this operant task within two weeks (Figure 1D). The most remarkable difference between groups was in task performance. Rats in the pre-training group manipulated the pedals randomly whereas rats in the post-training group chose the appropriate pedals and manipulated them based on the task rule (proportion of correct choices [%], pre-training: 51.0 ± 3.6; post-: 82.9 ± 1.8; Mann–Whitney test, *z* = 4.13, *p* < 3.6 × 10^−5^, *r* = 0.83; Figures 1D and S1C).

### Recording and cell-type classification in hippocampal formation

We recorded multineuronal spike activity i.e., multiple isolated single units, through silicon probes in the hippocampal formation (CA1 and LEC) of rats performing the self-paced, spontaneous left or right pedal pressing task. We first verified the connectivity patterns of LEC using two retrograde tracers, Fluoro-Gold and mRFP expressing glycoprotein-deleted rabies viral vector (ΔG-RV, Ohara et al., 2013). In line with previous studies (Witter et al., 2017; Ohara et al., 2018), injection of Fluoro-Gold into mPFC generated labeled neurons in layer Va, while ΔG-RV injection into DG and CA1 generated mRFP-labeled neurons mainly in layer IIa and III of LEC (Figure 2A). In this study, we focused on mPFC- and CA1-projecting LEC neurons which are found mostly in layer Va and III, respectively, which enabled us to increase the certainty of reconstructing the layer position of recorded neurons. We identified these neuron classes using the Multi-Linc method (Saiki et al., 2018; Soma et al., 2017; Nonomura et al., 2018; Rios et al., 2019) with antidromic stimulation of the ipsilateral mPFC for mPFC-projecting neurons and of the ipsilateral CA1 for CA1-projecting neurons (Figure 2B and C). We reconstructed the recording and stimulating sites histologically after recording (Figure 2B).

**Figure 2.**
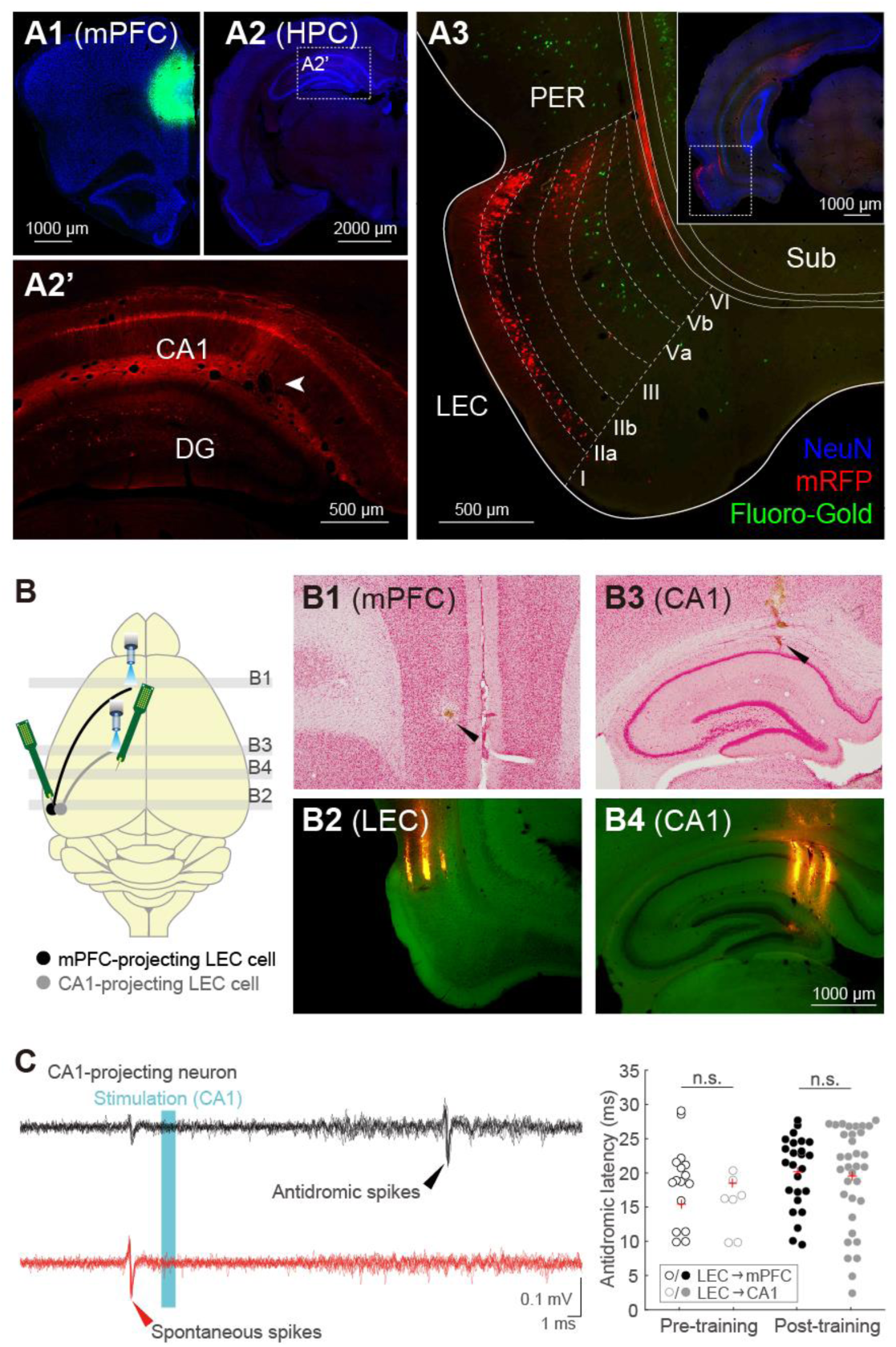
Efficient LEC recording with Multi-Linc methods based on the unique distribution of CA1-projecting LEC superficial neurons. (A) LEC neurons projecting to mPFC and HPC. Retrograde tracer (Fluoro-Gold) was injected into mPFC (A1) while retrograde viral tracer (mRFP-expressing G-deleted rabies viral vector) was injected into HPC, and the distribution of retrogradely labeled neurons was examined in LEC (A3). Note that mPFC-projecting neurons (green) distribute in layer Va while HPC-projecting neurons (red) distribute in layers IIa and III of LEC. (B) Schema showing the position of optical fibers for identifying the two different projection neurons in the LEC (left). The ipsilateral medial prefrontal cortex (mPFC) and CA1 were stimulated to identify mPFC- and CA1-projecting LEC neurons, respectively. Top-right images show the stimulation sites in mPFC (B1) and CA1 (B3). Track of the optical fibers (arrowhead) into the mPFC and CA1 in the Nissl-stained sections. Right-bottom images show recording sites in LEC (B2) and CA1 (B4). Probe shank tracks were visualized by fluorescent DiI (red). (C) Example of recordings from CA1-projecting LEC neurons during optical stimulation (cyan area), with spike collisions. Black and red traces represent antidromic spikes to optical stimulation and spike collision tests, respectively. Black arrowhead indicates antidromic spikes. Red arrowhead indicates the precedence of spontaneous spikes used as triggers for optical stimulation in collision tests (left). Spike latency after antidromic stimulation in mPFC- (black) and CA1-projecting (gray) LEC neurons (right).

The left panel of Figure 2C shows typical traces of antidromic spikes (black) and their disappearance due to collision with spontaneous spikes (red) in CA1-projecting LEC neurons. Since LEC neurons near the rhinal fissure project into the dorsal CA1 (Figure 2A), we could verify our recording electrode located in the LEC based on the presence of such CA1-projecting neurons (see also Figure 7), resulting in efficient *in vivo* LEC recording of behaving rats (Figures 2B and S2). We compared the antidromic latency between CA1- and mPFC-projecting neurons and found no difference (pre-training: LEC → mPFC cells, *n* = 7, mean ± SD, 19.3 ± 5.8 ms, LEC → CA1 cells, *n* = 16, 16.3 ± 4.1 ms; Mann–Whitney test, *z* = −1.44, *p* = 0.15, *r* = 0.30; Post: LEC → mPFC cells, *n* = 25, 21.0 ± 5.2 ms, LEC → CA1 cells, *n* = 34, 20.4 ± 7.3 ms; *z* = 0.00, *p* = 0.99, *r* = 0.00).

We isolated a total of 829 CA1 neurons and 1,287 LEC neurons from our multineuronal recordings during task performance. These neurons were further classified as either regular-spiking (RS, mostly putative excitatory neurons) or fast-spiking (FS, putative inhibitory neurons) neurons based on the bimodality of spike duration (Figure S2). Consistent with previous reports (Csicsvari et al., 1999; Nilssen et al., 2018), the ongoing spike rates of FS subtypes were significantly higher than those of RS subtypes in the CA1 compared to LEC (Figure S2). Given the small sample size of task-related FS neurons, we used RS neurons for further analysis.

### Development of task-related activities in CA1 and LEC cells

We observed the various types of task-related neurons in the CA1 and LEC of rats performing the self-paced forelimb pedal pressing task. We classified task-related neurons as Hold-type, Hold+Reward-type, Go-type, Go+Reward-type, and Reward-type using their preferred activities, namely contralateral or ipsilateral trials (Figures 3 and S3; see Methods), with task relevance indices based on previous studies (Soma et al., 2017, 2019; Rios et al., 2019). Figure 3A shows the peri-event time histograms (PETHs) of representative neurons which were related to pedal release (Go-type in the CA1, Figure 3A, left), to reward delivery and/or consume (Reward-type in the LEC, Figure 3A, middle), and to both pedal release and reward (Go+Reward-type in the LEC, Figure 3A, right; see also Figures 4-6 and S3). In the pre-training group, there were few task-related cells in the CA1 and LEC. In contrast, the fractions of task-related cells were dramatically increased after the completion of the task rule in both CA1 and LEC (Figure 3B; CA1-RS: pre-training, task-related cells, *n* = 22, non-task-related cells, *n* = 225; post-training, task-related cells, *n* = 195, non-task-related cells, *n* = 238; χ^2^ test, χ^2^ = 94.5, *p* < 2.5 × 10^−22^, *φ* = 0.37; LEC-RS: pre-training, task-related cells, *n* = 2, non-task-related cells, *n* = 333; post-training, task-related cells, *n* = 195, non-task-related cells, *n* = 594; χ^2^ = 94.6, *p* < 2.3 × 10^−22^, *φ* = 0.53).

**Figure 3.**
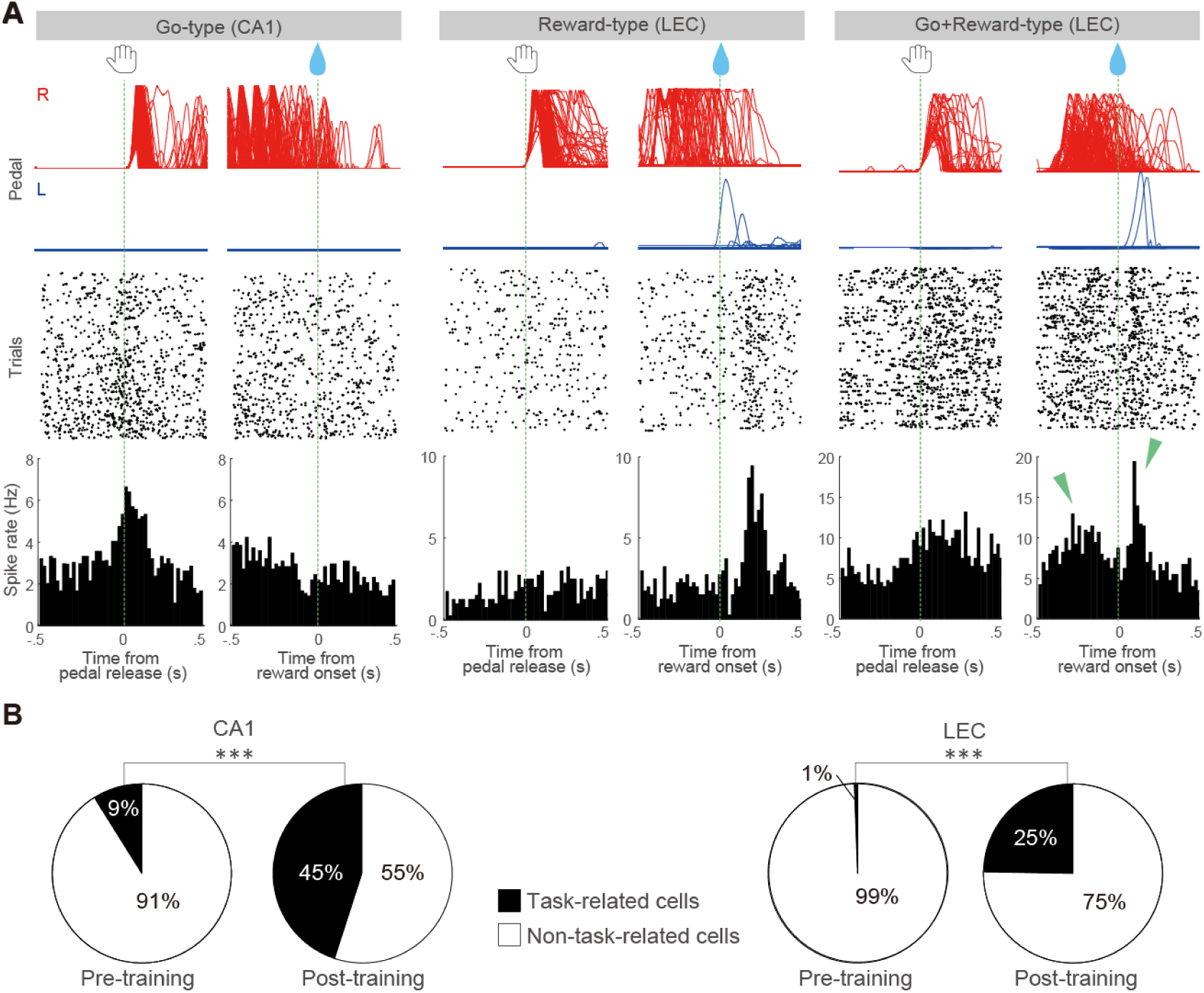
Task-related activities were dramatically developed after learning. (A) Examples of Go-type (left), Reward-type (middle), and Go+Reward-type (right) task-related activities in the CA1 (left) and LEC (middle, right) after learning. Go+Reward-type neuron shows increased activities in both pedal release and reward delivery periods (two peaks indicated by green arrowheads). Top, middle, and bottom show pedal trajectories (red: right pedal, blue: left pedal), spike raster plots, and peri-event time histograms (PETHs, Bin width, 20 ms), respectively. Spike data were aligned with the pedal release onset (left) or reward onset (right) at 0 s for individual task-related neurons. (B) Population ratios of task-related neurons in the CA1 and LEC of pre- and post-training groups. *** *p* < 0.001, 2 × 2 χ^2^ test.

**Figure 4.**
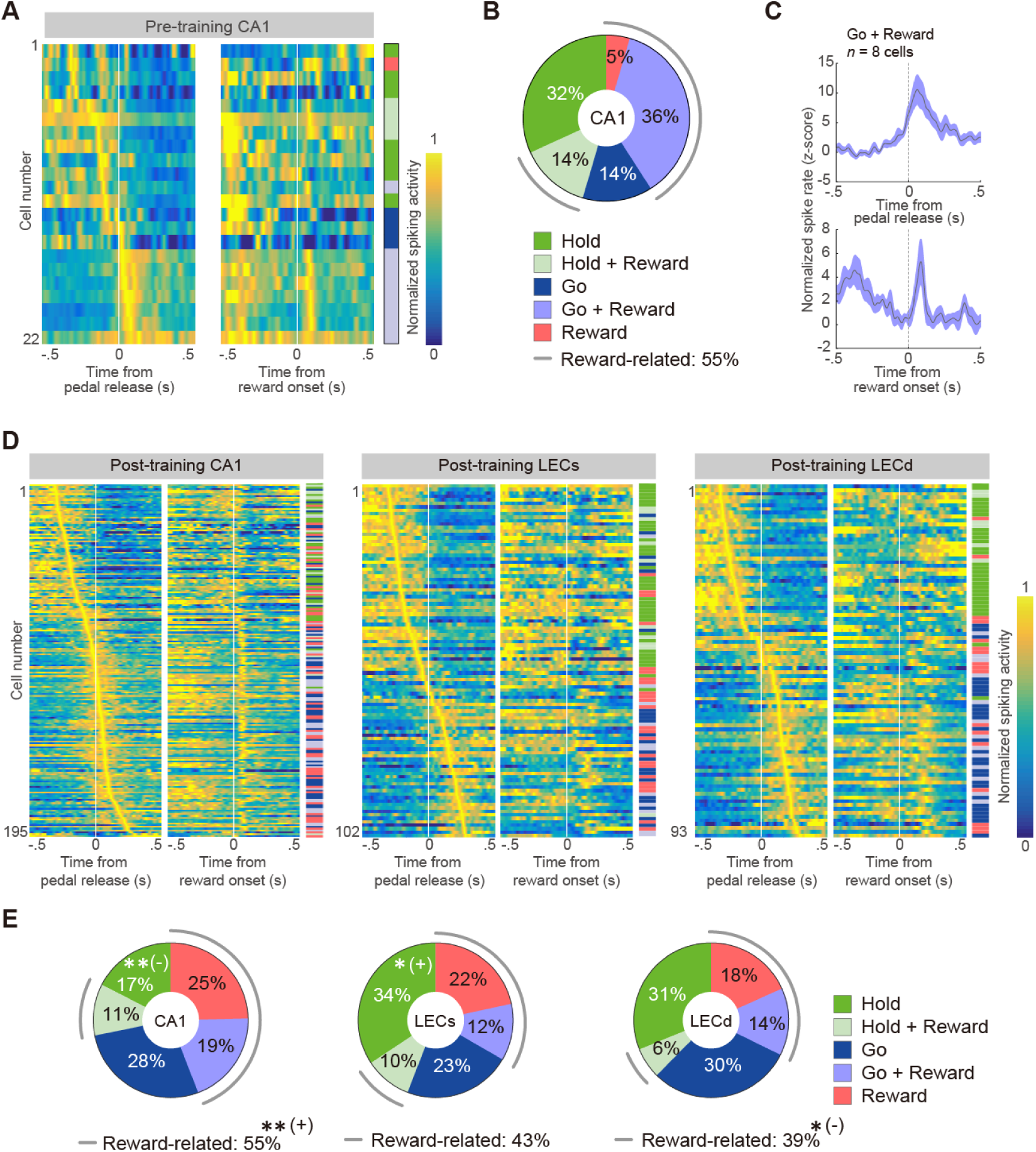
The repertoire of task-related activities in CA1 neurons before and after learning. (A) Five types of task-related activities in the RS subtype of CA1 neurons before learning. Normalized Gaussian-filtered PETHs (s = 12.5 ms for spikes in 0.05 ms bins) aligned with pedal release onset and reward onset at 0 s (vertical line) for individual task-related neurons. Each row represents a single neuron; they were sorted by the order of peak time obtained from pedal release onset data (early to late). The task-related type is indicated on the right side. (B) Population ratios of task-related types in CA1-RS neurons. (C) Averaged PETHs of all Go+Reward-type activities in CA1. PETHs were aligned with the pedal release onset (top) and reward onset at 0 s (bottom). Shaded regions represent 95% confidence intervals (CIs). (D) Five types of task-related activities in the RS subtypes of CA1 and LEC after learning. The figure legend is the same as in Figure 4A. (E) Population ratios of task-related types in the RS neurons of CA1 and LEC after learning. The figure legend is the same as in Figure 4B. ** *p* < 0.01, Residual analysis after 5 × 2 χ^2^ test.

In the pre-training group, CA1 neurons included five types of task-related neurons (Figure 4A and B; CA1-RS: Hold-type, *n* = 7; Hold + Reward-, *n* = 3; Go-, *n* = 3; Go +Reward-, *n* = 8; Reward-, *n* = 1). We found that the dominant fraction of task-related neurons represented both pedal release and reward (Go+Reward-type) before task learning (Figure 4B and C). In contrast, only two LEC neurons represented the task-related activity (LEC-RS: Hold-type, *n* = 0; Hold+Reward-, *n* = 0; Go-, *n* = 1; Go+Reward-, *n* = 0; Reward-, *n* = 1).

Next, we determined the types of task-related activity that were developed after learning in the CA1 and LEC (Figure 4D and E). Since the LEC neurons in the superficial and deep layers send their projections to different brain regions (Witter et al., 2017), we grouped the putative superficial layer cells (LECs) and putative deep layer cells (LECd) based on the results of Multi-Linc and histological observations (Figure 2). After learning, all five types of task-related activities were developed in both CA1 and LEC (Figure 4D; CA1-RS: Hold-type, n = 34; Hold+Reward-, *n* = 21; Go-, *n* = 54; Go+Reward-, *n* = 38; Reward-, *n* = 48; LECs-RS: Hold-, *n* = 35; Hold+Reward-, n = 10; Go-, n = 23; Go+Reward-, *n* = 12; Reward-, *n* = 22; LECd-RS: Hold-, *n* = 29; Hold+Reward-, *n* = 6; Go-, *n* = 28; Go+Reward-, *n* = 13; Reward-, *n* = 17). LECs and CA1 had significantly larger and smaller populations of Hold-type, respectively (Figure 4E; χ^2^ test, χ^2^ = 15.7, *p* < 4.7 × 10^−2^, *φ* = 0.20; post hoc residual analysis, CA1, *p* < 0.05; LECs *p* < 0.01).

Since the outcome-related activities seemed dominant in CA1 compared to LEC, we conducted further analysis after pooling the reward-related activity types as Reward-related type (CA1-RS: *n* = 107 [54.9%], LECs-RS: n = 44 [43.1%], LECd-RS: *n* = 36 [38.7%]). As expected, outcome-related information was predominantly encoded by CA1 (χ^2^ test, χ^2^ = 7.9, *p* < 2.0 × 10^−2^, *φ* = 0.14; post hoc residual analysis, *p* < 0.01). Also, LECd showed a significantly smaller population of outcome-related type than others (*p* < 0.05). Thus, both CA1 and LEC neurons came to show the various repertoire of task-related activities after learning.

### Temporal dynamics of action- and outcome-related CA1 and LEC neurons after learning

To visualize the temporal dynamics of the task-related neurons, we calculated averaged PETHs (Figure 5). Hold- and Hold+Reward-type neurons did not have a clear peak whereas others (Go-, Go+Reward-, and Reward-type neurons) showed clear peaks in both CA1 and LEC. At the population level, CA1 neurons had a shorter peak latency than LEC neurons (arrows, Figure 5). To quantify this observation, we calculated peak latency for both CA1 and LEC (Figure 6). First, we compared the peak latency of Go- and Go+Reward-types obtained from the PETHs aligned with pedal released and found that CA1 had a significantly shorter latency than LEC for both activity types (Go-type: CA1, −5.6 ± 133.9 ms; LECs, 72.3 ± 188.7 ms; LECd, 151.3 ± 97.4 ms; Kruskal–Wallis test, χ^2^ = 29.7, *p* < 3.6 × 10^−7^, *η*^*2*^= 0.23; post hoc Steel-Dwass test, CA1 vs. LECs, *p* < 3.6 × 10^−7^, *t* = 2.95, Cliff’s *d* = 0.43; CA1 vs. LECd, *p* < 6.1 × 10^−7^, *t* = 5.41, Cliff’s *d* = 0.73; LECs vs. LECd, *p* = 0.15, *t* = 1.04, Cliff ‘s *d* = 0.17, Figure 6A, top; Go+Reward-type: CA1, 55.9 ± 103.1 ms; LECs, 136.0 ± 151.1 ms; LECd, 128.7 ± 91.3 ms; χ^2^ = 8.4, *p* < 1.5 × 10^−2^, *η*^*2*^= 0.12; CA1 vs. LECs, *p* < 1.2 × 10^−2^, *t* = 2.39, Cliff’s *d* = 0.46; CA1 vs. LECd, *p* < 1.9 × 10^−2^, *t* = 2.13, Cliff’s *d* = 0.40; LECs vs. LECd, *p* = 0.23, *t* = 0.70, Cliff’s *d* = −0.17, Figure 6A, bottom). We also compared this parameter for Reward- and Go+Reward-types obtained from PETHs aligned with reward delivery. As for Go+Reward-type, there was no significant difference in peak latency among the three groups (Go+Reward-type: CA1, 78.5 ± 67.1 ms; LECs, 165.3 ± 87.6 ms; LECd, 182.8 ± 57.7 ms; χ^2^ = 25.1, *p* < 3.6 × 10^−6^, *η*^*2*^= 0.23; CA1 vs. LECs, *p* < 1.2 × 10^−2^, *t* = 2.39, Cliff’s *d* = 0.46; CA1 vs. LECd, *p* < 1.9 × 10^−2^, *t* = 2.13, Cliff’s *d* = 0.40; LECs vs. LECd, *p* = 0.23, *t* = 0.70, Cliff’s *d* = −0.17, Figure 6B, top). Conversely, Reward-type CA1 neurons showed a significantly shorter peak latency than LEC neurons in both superficial and deep layers (Reward-type: CA1, 93.4 ± 74.1 ms; LECs, 93.3 ± 94.4 ms; LECd, 126.4 ± 78.6 ms; Kruskal–Wallis test, χ^2^ = 2.2, *p* = 0.34, *η*^*2*^= 0.04; post hoc Steel-Dwass test, CA1 vs. LECs, *p* = 0.23, *t* = 0.80, Cliff’s *d* = −0.15; CA1 vs. LECd, *p* = 0.09, *t* = 1.32, Cliff’s *d* = 0.25; LECs vs. LECd, *p* = 0.18, *t* = 0.98, Cliff’s *d* = 0.23, Figure 6B, bottom).

**Figure 5.**
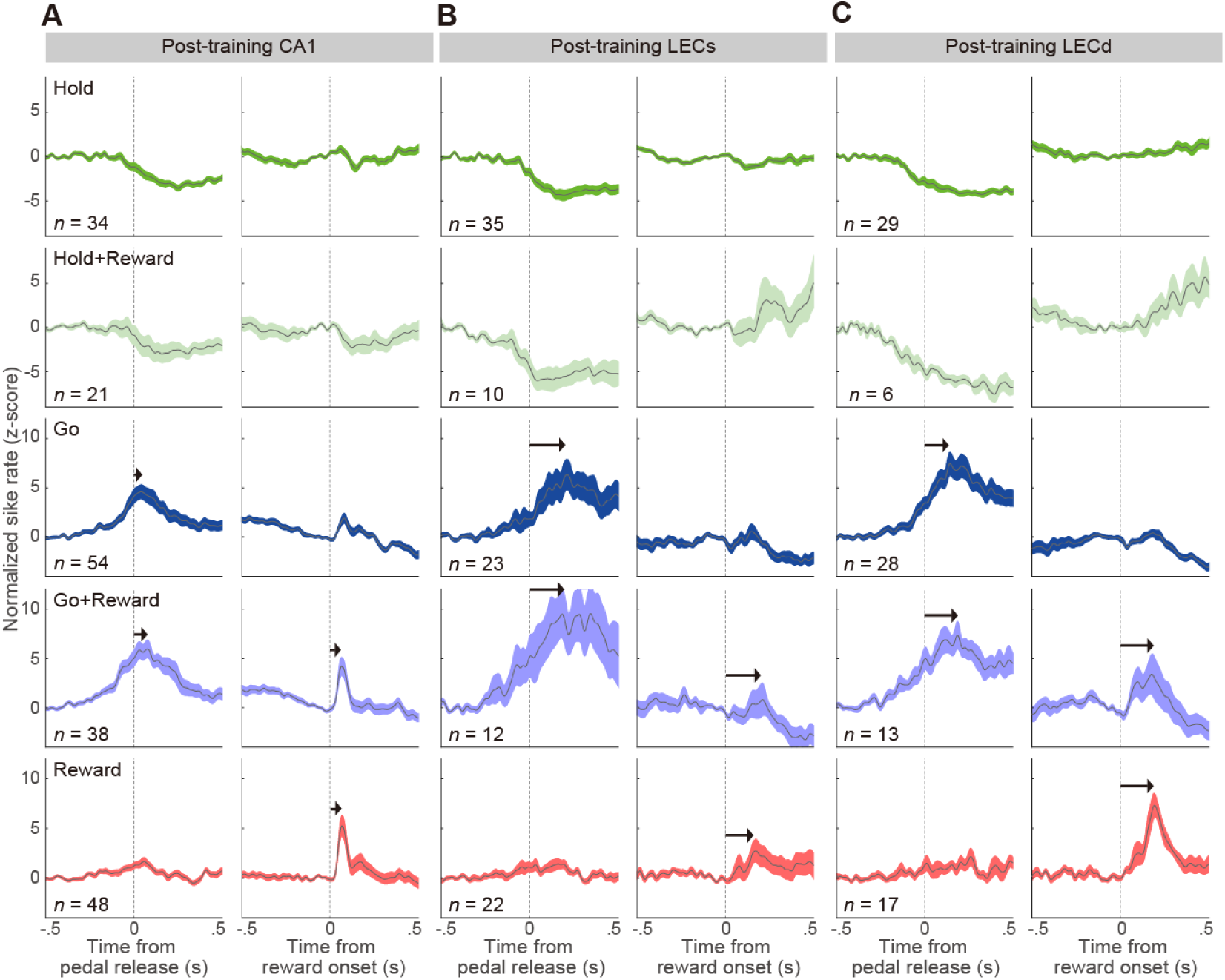
Population data for different types of task-related activities in the CA1 and LEC. (A-C) Averaged PETHs of all Hold-, Hold+Reward-, Go-, Go+Reward-, Reward-type activities in the CA1 (A), superficial (B), and deep layers of the LEC (C). PETHs were aligned with the pedal release onset (left columns) and reward onset at 0 s (right columns). Shaded regions represent 95% CIs. Arrows show the peak time.

**Figure 6.**
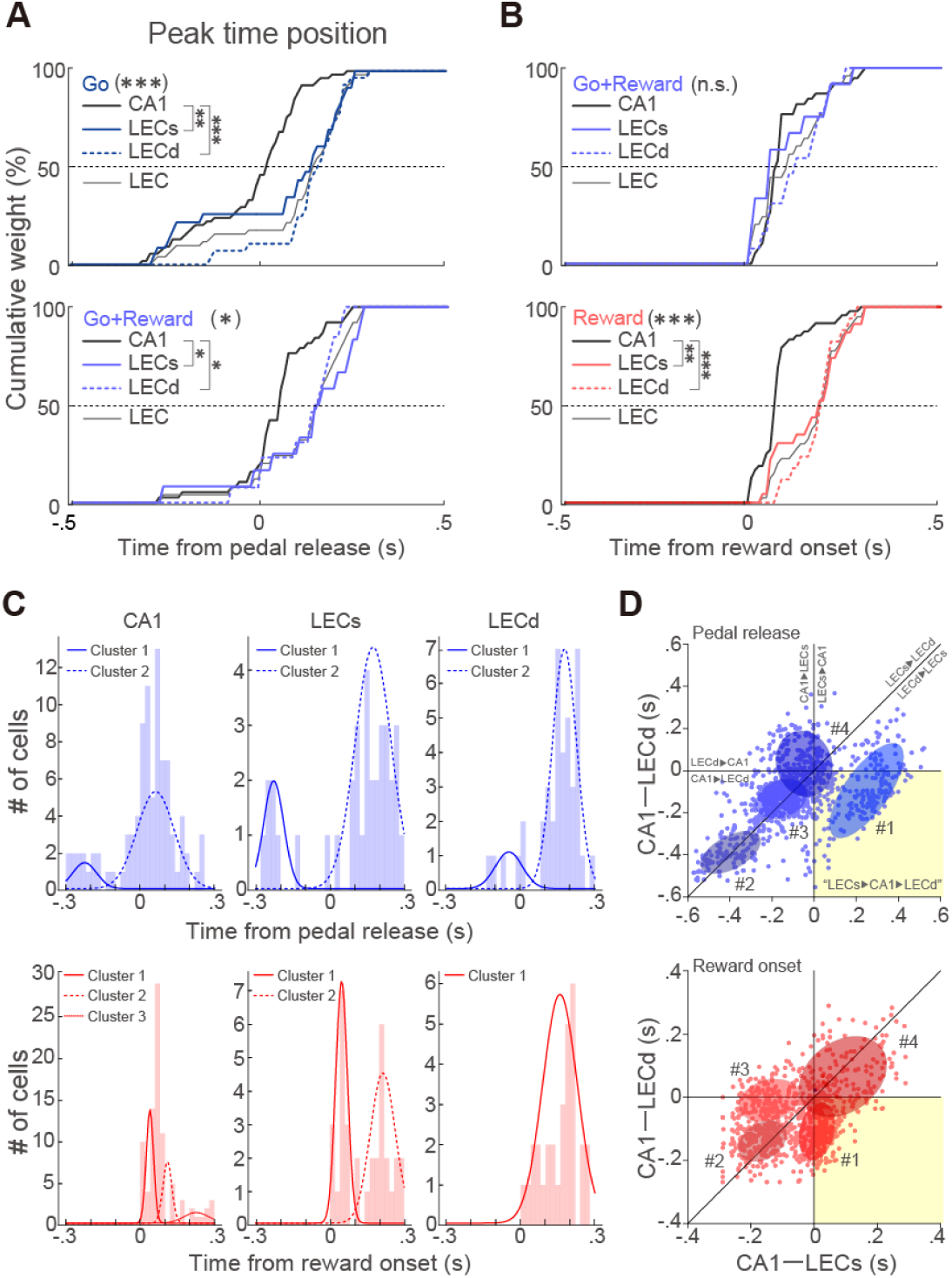
Comparison of peak time among CA1, superficial and deep LEC layers. (A and B) Cumulative distributions of peak time position from the onset of pedal release [A; Go-(top) and Go+Reward-type (bottom)] and that of reward delivery [B; Go+Reward-(top) and Reward-type (bottom)]. LECs and LECd represent the superficial and deep LEC layers, respectively. * *p* < 0.05, ** *p* < 0.01, *** *p* < 0.001, post hoc Steel-Dwass test. (C) Distribution of peak time positions from the onset of pedal release (top) and reward delivery (bottom) in the CA1 (left) as well as superficial (middle) and deep (right) LEC layers. The number of clusters was determined based on Bayesian information criterion (BIC) for the results of the Gaussian mixture model (GMM) fitting with the expectation-maximization (EM) algorithm (see Methods and Figure S4). (D) A bootstrap analysis (1000 samples) was performed on data to visualize the possible information flows. The number of clusters was determined based on the BIC for the results of the 2D GMM fitting with the EM algorithm (see Methods). Horizontal, vertical, and diagonal lines indicate borders of firing orders of the three regions (see appended annotations). Pale yellow background indicates a quadrant correspond to what one would expect based on the ECs → CA1 → ECd circuits.

The shape of the cumulative curves seems to comprise several ramps, which suggests the presence of subpopulations with different peak time positions. We tested this point by fitting a peak latency histogram to the Gaussian mixture model (GMM; Figure 6C). Cluster numbers were determined based on the Bayesian information criterion (BIC; Figure S4A). As for action-related activity types (Go- and Go+Reward-types), all three regions showed two distinct subpopulations: those preceding and those following spiking activity relative to pedal-release (Figure 6C, top CA1; mean ± SD in ms, −224.7 ± 48.7 and 62.2 ± 75.1; LECs, −227.4 ± 40.8 and 174.8 ± 72.8; LECd, −48.0 ± 54.7 and 177.0 ± 49.0). We verified this result with another method, *x*-means clustering (Figure S4B, top). As for outcome-related activity types, the latencies relative to the reward onset timing were clustered into multiple groups in some regions (CA1 = 3 cluster, 39.2 ± 14.8, 110.0 ± 18.1, and 226.7 ± 50.2; LECs = 2 clusters; 47.3 ± 23.7 and 214.2 ± 50.8; LECd = 1 cluster, 158.7 ± 70.2, mean ± SD in ms; Figure 6C, bottom) although *x*-means clustering showed that all three regions had two distinct subpopulations (Figure S4B, bottom). These results enabled us to speculate that distinct subpopulations send their signals to others with different timings.

To visualize pseudo-signal flow among the three regions, we calculated the pseudo-paired differences of peak latency between two-different area neurons (i.e., CA1 vs LECs and CA1 vs. LECd; Figures 6D and S4C; see Methods). We found that action-related neurons included the subpopulation reflecting the hippocampal-entorhinal circuit (cluster #1 in the pale-yellow background, LECs → CA1 → LECd; Figure 6D, top) and other clusters (#2 and 3) showed that the CA1 neurons act before LEC neurons. Instead of a subpopulation reflecting the hippocampal-entorhinal circuit, the outcome-related neurons included other types of subpopulations showing that CA1 and LECs neurons act simultaneously and send their signals to LECd neurons (cluster #1; Figure 6D, bottom). As shown in Figures 5 and 6B, cluster #2 showed that CA1 neurons act prior to LEC neurons. These results suggest the existence of subpopulations of CA1 and LEC neurons that process information in a different order from that suggested by previous anatomical findings i.e., the hippocampal-entorhinal circuit.

### Firing rate and limb specificity of action- and outcome-related CA1 and LEC neurons after learning

We compared the firing rate of hold-related activities (Hold- and Hold+Reward-type) by averaging the spike activities during the pre-movement period (−1000 to −500 m). LECd showed a significantly lower firing rate than both CA1 and LECs (Figure S5A and B, top). As for action-related (Go- and Go+Reward-type) and outcome-related (Reward- and Go+Reward-type) activities, we calculated peak activities (peak ± 150 ms). In the action-related types, CA1 showed significantly higher spiking activities than both LECs and LECd (Figure S5A and B, middle). Additionally, the peak activities of outcome-related types of CA1 were significantly higher than those of LEC. In addition, significant difference in peak activities was observed between LECs and LECd (Figure S5A and B, bottom). Thus, we found contrasting peak activities in superficial versus deep LEC layers.

Since our original task can evaluate the laterality i.e. the preference for contra- or ipsilateral limb movement of task-related activity, we tested if CA1 and LEC neurons show lateralized activity. As for action-related activities (Go- and Go+Reward-types), Go-type CA1 neurons preferred ipsilateral activity but other subpopulations did not show lateralized activities (Table S1). Thus, both CA1 and LEC neurons have basically bilateral activity, and the ipsilateral preference of Go-type CA1 neurons is similar to that of the posterior parietal cortex (PPC; Soma et al., 2019). As for outcome-related activities (Go+Reward- and Reward-types), both CA1 and LEC did not show evidence of laterality (Table S2).

### Task-related activities of mPFC- and CA1-projecting neurons in the LEC

Finally, we examined task-related activities of identified projection LEC neurons (Figure 7). Figure 7A is a representative example of an mPFC-projecting LEC neuron that showed Go-type activity, which suggests that this neuron sends action-related information to the mPFC. Figure 7B shows the task-related activities of all identified neurons. Consistent with the current (Figure 2A) and previous histological observations (Ohara et al., 2018), the mPFC- and CA1-projecting LEC neurons were recorded from deep and superficial layers, respectively. The only exception was mPFC-projecting LEC neurons in the superficial layer which likely corresponds to calbindin-positive neurons in LEC layer IIb (Ohara et al., 2019). Both projection neurons showed action- and outcome-related activities. The outcome-related activities were often observed in CA1-projecting LEC neurons rather than mPFC-projecting neurons (LEC → dCA1: 6/10 cells [60%]; LEC → mPFC: 3/9 cells [33.3%]). Although this tendency was not tested for statistical significance because of the small sample size, we obtained a similar result by comparing reward-related fractions between LECd and LECs (Figure 4E). These data indicate that the LEC sends the internal context (action and outcome information) to the CA1 and mPFC after learning.

**Figure 7.**
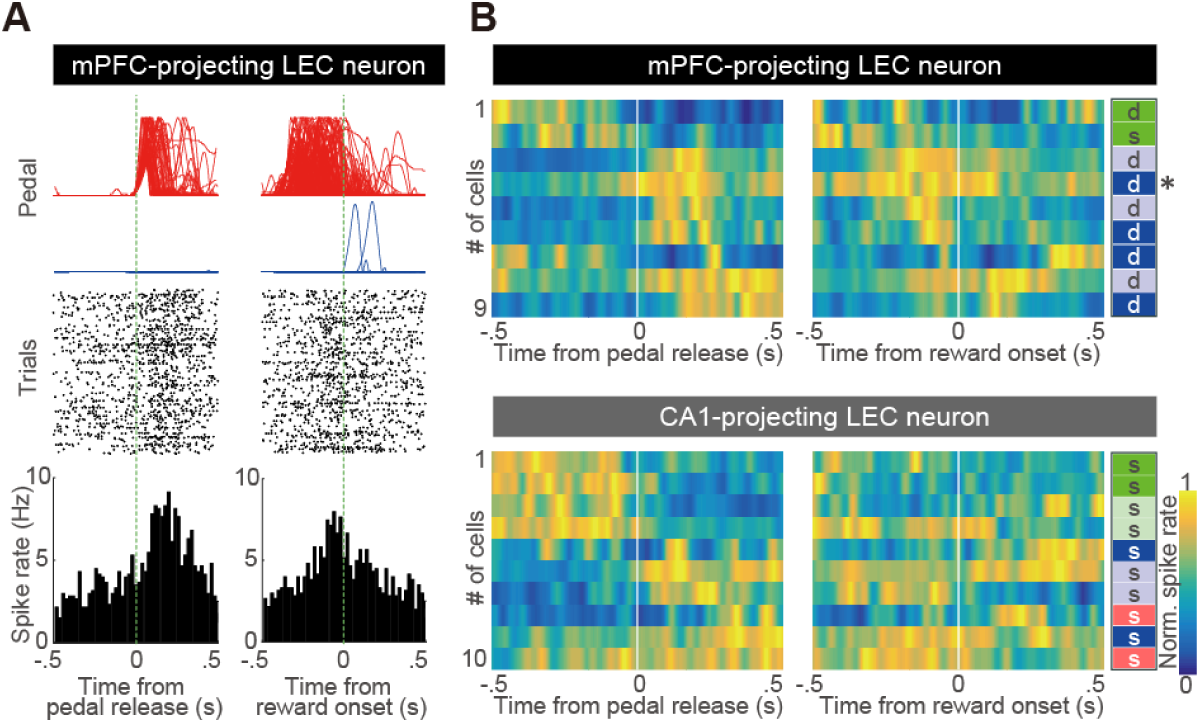
Task-related activity of mPFC- and CA1-projecting neurons. (A) Examples of Go-type task-related activity in the LEC. This neuron was identified as an mPFC-projecting neuron with Multi-Linc methods (see Methods). The figure legend is the same as Figure 3A. (B) Task-related activity of mPFC-(top) and CA1-projecting (bottom) neurons in the LEC. The figure legend is the same as Figure 4A. The task-related type is indicated on the right side with layer position (s: superficial layer, d: deep layer). * represents the example neuron shown in Figure 7A.

## Discussion

To investigate the timing of neural representation for two distinct behavioral events related to learning in the hippocampal-entorhinal circuit, we recorded neuronal activities in CA1 and LEC (superficial and deep layers) neurons while rats performed the simplest operant learning task requiring a spontaneous action of pedal release to acquire a reward. We made it possible to monitor the timing of distinct behavioral events involving action and reward precisely by using head-fixation instead of the standard freely-moving condition (Figures 1 and S1; Soma et al., 2017; 2019). Also, we presumed the layers of the recording site in LEC, either superficial or deep layers, by optogenetically identifying mPFC- and CA1-projecting LEC neurons based on anatomical findings that superficial CA1-projecting LEC neurons were locally distributed around the dorsal border of LEC (Figure 2; Ohara et al., 2018). These experimental setups enabled us to distinguish the types of task-related activities related to action and reward events and precisely compare the timing of these activities for CA1 neurons and for superficial and deep layer LEC neurons in the hippocampal-entorhinal circuit.

In summary, our main findings are: (1) the ratio of task-related neurons that showed task-related activity increased in both CA1 and LEC after learning (Figure 3); (2) five types of task-related activities (Go-type, Go+Reward-type, Hold-type, Hold+Reward-type, and Reward-type) were observed and the LEC developed a large population of the Hold-type compared to CA1 (Figure 4); (3) peak latency was shorter for CA1 than LEC among Go-type, Go+Reward-type, and Reward-type activities (Figures 5, 6A, and 6B); (4) each area had distinct clusters showing different peak time positions (Figure 6C) and action-related neurons included subpopulations reflecting the hippocampal-entorhinal circuit, whereas the outcome-related ones did not contain such subpopulations (Figure 6D); and (5) the mPFC- and CA1-projecting LEC neurons identified with Multi-Linc (Figure 2) represented both action- and outcome-related information (Figure 7).

### The appearance of task-related activities after learning

Both CA1 and LEC neurons developed task-related activities after learning. These task-related activities in the hippocampal-entorhinal circuit were thought to be neuronal representations that were acquired while animals had experienced various episodic events during operant learning. Before the rats learned the task rule, only a few LEC neurons showed task-related activities (<1%); however, a substantial number of CA1 neurons responded to behavioral events (9%; Figure 3). These CA1 neurons mainly consist of the Go+Reward-type (Figure 4), which suggests that there are CA1 neurons (undifferentiated neurons) sensitive to multiple events in the early phase of learning. Moreover, these neurons contribute to the formation of action-outcome contingency through episodic events. In contrast, task-related LEC neurons appeared de novo after learning (Figure 3). LEC neurons can represent different types of information after learning (Deshmukh and Knierim, 2011; Deshmukh et al., 2012; Lu et al., 2013; Tsao et al., 2013; Igarashi et al., 2014; Keene et al., 2016; Li et al., 2016; Pilkiw et al., 2017; Tsao et al., 2018; Wang et al., 2018; Lee et al., 2021). These distinctions between CA1 and LEC neurons suggest that CA1 acts as a foundation for providing episodic information about operant learning to the LEC in the early phase of learning so that the LEC can adapt to handle this information as learning progresses (cf. Igarashi et al., 2014). In addition to the go- and reward-related types, neurons that exhibited hold-related activities appeared after learning, indicating that these activities were not simple waiting activities as previously observed in the motor cortex (Soma et al., 2017) but task-relevant ones which may represent the time to express the specific behavior involved in operant learning. In fact, Hold-type neurons were mostly distributed in the LEC (Figure 4). Additionally, relatively longer time representation has been reported in the LEC (Tsao et al., 2018).

### Distinct timing of action representation by CA1 versus LEC

Both CA1 and LEC neurons showed essentially bilateral preference although Go-type of CA1 neurons had a subtle ipsilateral bias (Table S1). These results indicate that go-related activity in the CA1 and LEC could represent abstract information for motion expression. Similar to these areas, previous studies demonstrated that the limb specificity of voluntary forelimb movement is bilateral or slightly biased to ipsilateral in the PPC, and PPC is involved in such abstract information whereas the primary motor cortex showed contralateral representation and related to concrete motor information (Soma et al., 2017, 2019). Peak time position of action-related types (Go- and Go+Reward-types) in the CA1 was shorter than that of LEC (Figures 5 and 6). Our analysis revealed two subpopulations in the CA1 and LEC: the preceding and following spiking activity relative to the pedal-release (Figure 6). The former is thought to be preparation or planning activity for voluntary movement and the latter could be feedback activities to regulate or monitor the expressing movements. In fact, the LEC directly received inputs from both PPC and motor cortices (Burwell and Amaral, 1998; Olsen et al., 2017). The visualization of pseudo-signal flow suggested that task-related information was processed though distinct subpopulations: LECs → CA1 → LECd or CA1 → LECs & LECd (Figure 6D, top). Thus, it revealed that CA1 and a portion of the LEC’s superficial layer neurons first represent the preparation or planning for voluntary expressing movement. Next, most of the remaining CA1 neurons act instantaneously, and the superficial and deep layer LEC neurons act in succession during actual movement in the hippocampal-entorhinal circuit.

### Distinct timing of reward representation by CA1 versus LEC

Based on anatomical studies, sensory information about the auditory signal for reward presentation (audition) and licking (somatosensation) is expected to be processed sequentially through the entorhinal-hippocampal-entorhinal pathway: LEC superficial layers → CA1 → LEC deep layers. In contrast to this expectation, reward-responsive activities (Reward-type and Go+Reward-type) of CA1 neurons preceded those of LEC neurons, and CA1 showed a sharp peak compared to the LEC (Figures 5 and 6). Since licking continues for a few seconds, it is unlikely that LEC neurons represent drinking behavior with licking. In the pre-training group, Reward-type neurons were rare in both CA1 and LEC (Figures 3 and 4). Therefore, it is likely that the Reward-type activity of CA1 and LEC neurons are not simple sensory responses but are learning-related activities that have developed through operant training.

A similar inconsistency between connectivity and activity patterns has also been reported by a study that examined the processing of olfactory sensory inputs using an in vitro–isolated guinea pig brain preparation (Biella and de Curtis, 2000). The neural activity induced by the lateral olfactory tract stimulation propagated sequentially from the LEC to the hippocampus, and from the hippocampus to the MEC but not the LEC. This finding, together with our results, implies that information processing through the entorhinal-hippocampal-entorhinal pathway is more complex than initially reported. One possible explanation for early reward representation in CA1 is the direct input of reward signals to CA1 that bypasses the EC. Indeed, CA1 receives direct dopaminergic inputs from the locus coeruleus (Takeuchi et al., 2016) as well as the ventral tegmental area (VTA). The superficial LEC also receives dopaminergic inputs from the VTA (Lee et al., 2021). Moreover, CA1 receives inputs from mPFC, via the nucleus reuniens of the thalamus, which are crucial for representing the future path during goal-directed behavior (Ito et al., 2015). This mPFC input to the CA1 may have contributed to the action-related activities of CA1 neurons which preceded those of LEC neurons. The intrahippocampal circuits, particularly recurrent circuits in the CA3 regions, may also play a role in amplifying event information from the LEC leading to the sharp activity peak observed in CA1.

### Significance of developed task-related activities in CA1 and EC

CA1 represented both the internal (voluntary action) and external (reward) events in contiguity with the actual timing of events i.e., in real-time compared with the LEC. It may be necessary for CA1 to form an episodic memory of *when-where-what* information. When an animal is at a certain time and place, time cells and place cells are activated in the CA1. In addition, if a certain event occurs, CA1 neurons immediately respond to that event, resulting in representation of the event as it occurs in real-time. Thus, CA1 could represent the specific event with specific spatiotemporal information i.e., by taking a snapshot of episodic (*when-where-what*) information. In contrast, the EC more universally represents spatiotemporal information (Fyhn et al., 2004; Hafting et al., 2005; Deshmukh et al., 2010; Tsao et al., 2018) in an ongoing manner possibly like a movie (Sugar and Moser, 2019). Triggered by the event, CA1 takes the snapshot from the entorhinal movie, and the EC processes the hippocampal snapshot in the space of universal information before transferring it to the mPFC as the central executive system. In this way, animals can use *when-where-what* information to optimize their behaviors through the entorhinal-hippocampal circuit.

## Methods

### Animals

All experiments were approved by the Animal Research Ethics Committee of Tamagawa University (animal experiment protocol, H22/27-32), and were carried out in accordance with the Fundamental Guidelines for Proper Conduct of Animal Experiment and Related Activities in Academic Research Institutions (Ministry of Education, Culture, Sports, Science, and Technology of Japan) and the Guidelines for Animal Experimentation in Neuroscience (Japan Neuroscience Society). All surgical procedures were performed under appropriate isoflurane anesthesia (see below). All effort was made to minimize suffering. The procedures for our animal experiments were established in our previous studies (Isomura et al., 2009; 2013; Kimura et al., 2017; Yoshida et al., 2018; Kawabata et al., 2020). This study is based on data from the channelrhodopsin-2 (ChR2)-expressing (Thy1-ChR2) transgenic rats (W-TChR2V4; *N* = 25 rats, male, 316 ± 39 g) abundantly expressing ChR2-Venus fusion protein under the control of the Thy1.2 promoter in cortical and other neurons (Tomita et al., 2009; Saiki et al., 2018). These animals were kept in their home cage under an inverted light schedule (lights off at 9 a.m., lights on at 9 p.m.).

### Surgery

Rats were handled briefly by the experimenter (10 min, twice) before the day of surgery. For head plate implantation, rats were anesthetized with isoflurane (4.5% for induction and 2.0–2.5% for maintenance; Pfizer Japan Inc., Tokyo, Japan) using an inhalation anesthesia apparatus (Univentor 400 anesthesia unit, Univentor, Zejtun, Malta) and placed on a stereotaxic frame (SR-10R-HT, Narishige, Tokyo, Japan). In addition, lidocaine jelly (AstraZeneca, Osaka, Japan) was administered around surgical incisions for local anesthesia. During anesthesia, body temperature was maintained at 37°C using an animal warmer (BWT-100, Bio Research Center, Tokyo, Japan). The head plate (CFR-2, Narishige) was attached to the skull with small anchor screws and dental resin cement (Super-Bond C&B, Sun Medical, Shiga, Japan; Unifast II, GC Corporation, Tokyo, Japan). Reference and ground electrodes (Teflon-coated silver wires, A-M systems, WA, USA; 125 µm in diameter) were implanted above the cerebellum. Analgesics and antibiotics were applied after the operation (meloxicam, 1 mg/kg s.c., Boehringer Ingelheim Japan, Tokyo, Japan; gentamicin ointment, 0.1% ad usum externum, MSD, Tokyo, Japan).

Water deprivation was started after full recovery from surgery (6 d later). The rats had *ad libitum* access to water during weekends but obtained water only by performing the task correctly during the rest of the week. When necessary, an agar block (containing 15 ml water) was given to the rats in their home cage to maintain them at >85% of their original body weight (Saiki et al., 2014; Soma et al., 2018, 2021).

### Behavioral task

We used the self-paced, spontaneous left or right pedal pressing task in our operant conditioning system (custom-made by O’hara & Co., Ltd., Tokyo, Japan; Fig. 1*A*; see also Soma et al., 2017; 2019; Rios et al., 2019) to examine the timing of neural representation of two distinct behavioral events related to learning in the CA1 and LEC. In this task, the rats had to manipulate left and right pedals with the corresponding forelimb in a head-fixed condition. They spontaneously started each trial by pushing both pedals down with both forelimbs and holding them down for a short period (“holding period,” at least 1 s; within a holding area, 0–30% in relative pedal position). After completing the holding period, the rats had to release either the left or the right pedal, depending on the context without any instruction cue, to obtain 0.1% saccharin water (10 μl) as a reward. The reward was dispensed from the tip of a spout by a micropump with a 300–700 ms delay (100 ms steps at random). This task consisted of two blocks of right-rewarded trials and left-rewarded trials. Each block lasted until the rat performed more than 30 correct (rewarded) trials and achieved 80% correct performance in the 10 most recent trials or or 100 rewards had been obtained. If the rats incorrectly released the other pedal (error trial) or failed to complete the holding period (immature trial), then they did not receive feedback. The rats typically learned this operant task within two weeks (2–3 h per day).

Once the rats completed task learning, they underwent a second surgery under isoflurane anesthesia for later recording experiment. We made tiny holes (1.0–1.5 mm in diameter) in the skull and dura mater above the CA1 (3.0 and 4.5 mm posterior, ± 2.0 mm lateral from bregma), LEC (6.0 mm posterior, ± 6.8 mm lateral), and the medial prefrontal cortex (mPFC; 3.5 mm anterior, ± 0.6 mm lateral). LEC and CA1 coordinates were determined by our previous study (Soma et al., 2013; Ohara et al., 2018). All holes were immediately covered with silicon sealant (DentSilicone-V, Shofu, Kyoto, Japan) until recording experiment.

### In vivo electrophysiological recording

We performed extracellular multi-neuronal (multiple isolated single-unit) recordings from individual neurons while the rats were performing behavioral tasks. Supported by agarose gel (2% agarose-HGT, Nacalai Tesque, Kyoto, Japan) on the brain, 32-channel silicon probes (Iso_3x_tet-A32 or Iso_4x_tet-A32; NeuroNexus Technologies, MI, USA; Saiki et al., 2018) were inserted into precisely the CA1 and LEC. Insertions were performed using fine micromanipulators (SM-15 or SMM-200B, Narishige) at least 1 h before the start of each recording experiment.

Wide-band signals were amplified and filtered (FA64I, Multi Channel Systems, Reutlingen, Germany; final gain, 2,000; band-pass filter, 0.5 Hz to 10 kHz) through a 32-channel head stage (MPA32I, Multi Channel Systems; gain, 10). These signals were digitized at 20 kHz and recorded with three 32-channel hard-disc recorders (LX-120, TEAC, Tokyo, Japan), which simultaneously digitized the pedal positions tracked by angle-encoders and the events of optogenetic stimulation.

### Optogenetic stimulation

We performed the Multi-Linc (multi-areal/multineuronal light-induced collision) method to effectively identify the pyramidal neurons sending direct projections to specific areas by combining multi-areal optogenetic stimulation and multi-neuronal recordings. Details of this procedure were described previously (Saiki et al., 2018). Briefly, prior to the insertion of silicon probes, the optical fibers (FT400EMT, FC, Thorlabs, NJ, USA; NA, 0.39; internal/external diameters, 400/425 μm) for stimulation were vertically inserted into the mPFC (4,100 μm deep) and CA1 (2,300 μm deep) using micromanipulators (SM-25A, Narishige). To evoke antidromic spikes in specific axonal projections from the LEC neurons (mPFC- and CA1-projecting cells), a blue LED light pulse (intensity, 5–10 mW; duration, 0.5–2 ms, typically 1 ms) was applied through each of the two optical fibers using an ultra-high-power LED light source (UHP-Mic-LED-460, FC, Prizmatix Ltd., Givat-Shmuel, Israel) and a stimulator (SEN-8203, Nihon Kohden, Tokyo, Japan). To be classified as projecting neurons, neurons were required to meet several criteria, including constant latency, fixed frequency (frequency-following test, two pulses at 100 and 200 Hz), and collision test (Lipski, 1981; Soma et al., 2017; Nonomura et al., 2018; Saiki et al., 2018; Rios et al., 2019).

### Spike isolation

Raw signal data were processed offline to isolate spike events of individual neurons in each tetrode of the silicon probes. Briefly, spike candidates were detected and clustered by our semiautomatic spike-sorting software, EToS (Takekawa et al., 2010, 2012). Using the clustering software Klusters and the viewing software NeuroScope (Hazan et al., 2006), spike clusters were manually combined, divided, discarded, or subject to a combination thereof to refine single-neuron clusters based on two criteria: the presence of refractory periods (>2 ms) in their own autocorrelograms and the absence of refractory periods in cross-correlograms with other clusters. We included single-neuron clusters if they had enough spike trains during task performance (≥20 trials with total ≥250 spikes). These clusters were classified as either task-related or non-task related neurons (Figure 3B; see also below).

### Spike collision analysis

To identify mPFC- and CA1-projecting LEC neurons, we used the Multi-Linc method with post hoc analysis to complete multi-neuronal collision tests (Saiki et al., 2018). Briefly, after offline sorting for spike isolation, we compared filtered traces with no spikes prior to the stimulus (control traces, black ones in Figure 2C) against those that had a spike in one spike cluster (test traces, red ones in Figure 2C) using MATLAB (MathWorks, MA, USA). If we found antidromic-like (all-or-none and no-jittering; black arrowheads in Figure 2C) spike activities with short latency in many of the control traces, we set a time window for counting possible antidromic spikes, based on a clear dissociation between averaged control and test traces due to presence or absence of spikes. The cut-off threshold defined in a receiver operating characteristic curve for distributing most-negative points (trough of spike waveform) within the time window was used to determine whether spikes were present, so that we obtained spike and no-spike counts in the control and test events. Based on this method, we included spike clusters with control spike probability above 50% and test spike probability less than half of the control. Finally, passing the collision test was statistically justified by a 2 × 2 χ^2^ test (*p* < 0.05) for spike and no-spike counts in control and test events (see Supplementary Figure 9 in Saiki et al., 2018). The latency of antidromic spikes was defined as the time from the onset of stimulation to the median of their peak positions within the time window, and their jitter was defined as the time between the first (25%) and the third (75%) quartiles of their peak positions within the time window. In this way, we judged if these spikes were antidromic based on collisional disappearance of antidromic spikes (collision test), as well as their all-or-none properties, absence of jitters (constant latency test; <0.5 ms), and high reliability (frequency-following test; if applicable in the tentative collision test).

### Spike analysis

Within each neuron (spike cluster), basal spiking properties and task-related activity in relation to behavioral task performance were analyzed using MATLAB as follows. The ongoing spike rate and spike duration (onset to peak) for individual spike clusters were defined as in our previous studies (Isomura et al., 2009; Soma et al., 2017; Saiki et al., 2018). Spike clusters were classified as regular-spiking (RS, mostly putative excitatory) neurons and fast-spiking (FS, putative inhibitory) neurons based on spike duration (≥0.6 ms for RS neurons, <0.6 ms for FS neurons). Since we refer to many groups of neurons (RS vs. FS; and CA1 vs. LEC), we used abbreviations for simplicity e.g., CA1-RS for RS neurons in CA1.

Next, we examined task-related spike activity correlated with self-initiated action or outcome (reward delivery). For action-related activity, we analyzed spike trains in relation to unilateral forelimb movements during task performance (≥20 trials with total ≥250 spikes), which were aligned with the onset (0 s) of pedal release (following ≥1 s holding time). Release onset was defined as follows: pedal release was first detected as the time when the pedal went outside the holding area (30% in relative pedal position). Next, onset was defined as the time when the pedal position exceeded 5% in relative pedal position before approximate pedal release. The task-related activity was defined by the task relevance index using the Kolmogorov–Smirnov (KS) test, as previously described (Kimura et al., 2017; Saiki et al., 2018; see also Figure 2A in Soma et al., 2017); we defined a task-related neuron as a neuron with a task relevance index (*p*) smaller than the criterion (*p* = 10^−6^) in contra- or ipsilateral pedal release trials. The preference activity (contra- or ipsilateral) was defined as the side with the smaller task relevance index. Task-related neurons were further classified into Hold-type and Go-type according to the peak time position of spike increase and the dependence on pedal holding time in the peri-event time histogram (PETH; 20 ms bins) on the preferred side (Soma et al., 2017, 2019; Rios et al., 2019). A Hold-type neuron has a sustained spike increase prior to the release onset (0 s) depending on the holding time. A Go-type neuron has a phasic spike increase independent of the holding time.

To check the limb preference of Go-type neurons, we compared the peak amplitude of contralateral and ipsilateral release trials. Peak amplitude was calculated by averaging the spike rate in the peak period (center of peak bin ± 150 ms), in which the peak bin was determined in the PETH of preferred release trials (contralateral or ipsilateral). We used the same peak period to calculate peak amplitude in non-preferred release trials. For Hold-type neurons, we compared the mean spike rate during the holding period (−1,000 to 0 ms) between contralateral and ipsilateral release trials. Moreover, we evaluated the limb preference (laterality) of Go-type neurons using the laterality index (ranging from −1 to +1; Soma et al., 2017, 2019; Rios et al., 2019) based on normalized peak activities as follows,

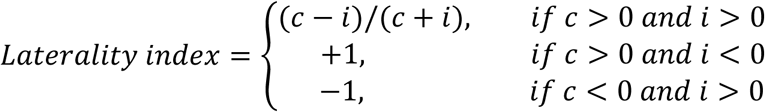

where *c* and *i* are activities associated with contralateral and ipsilateral movements, respectively. These parameters were obtained from the following equation:

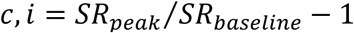

where *SR*_*peak*_ is the mean spike rate in the peak period (the center of peak bin ± 150 ms), and *SR*_*baseline*_ is the mean spike rate in the baseline period (−1,000 to −700 ms relative to pedal release onset). Consequently, laterality index values of −1 and +1 indicate ipsilateral- and contralateral-preferring neuronal activity, respectively.

For outcome-related activity, spike trains were aligned with the onset (0 s) of reward delivery (≥20 trials with total ≥250 spikes), and outcome-related activity (Reward-type) was defined in the same way as action-related activity. When neurons were classified as both action-related type and outcome-related type based on PETHs aligned with pedal release and reward delivery, these were called action- and outcome-related type, i.e., Go+Reward-type, or Hold+Reward-type, according to the action-related activity.

### Gaussian mixture model with Bayesian information criterion

Peak latencies extracted from PETHs of spike activity associated with pedal release and reward delivery were clustered into several groups based on the assumption of the Gaussian mixture model (GMM). We assumed that a distribution of the peak latencies could be represented with a small number of Gaussian distributions. Basically, the number of Gaussian distributions should be set by a user in advance as a hyper parameter. We repeated the GMM fitting with the expectation-maximization (EM) algorithm 1000 times for cluster numbers of 1 to 5, and calculated a Bayesian information criterion (BIC) for each repeat:

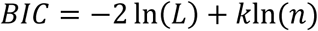

where *L, k*, and *n* indicate likelihood of each sample, number of parameters, and number of samples, respectively. Then we defined an optimal cluster number which showed a minimum mean BIC (Figure S4A). We used the *x*-means clustering algorithm as well (Pelleg and Moore, 2000) (Figure S4B), which is an extension of the *k*-means clustering algorithm, for re-confirming clustering results with an algorithm other than the GMM clustering algorithm.

For calculating pseudo-paired differences in peak latencies between two-different area neurons (i.e., CA1 vs LECs and CA1 vs. LECd), we conducted a 1000-repeat bootstrap analysis, and clustered these differences into several groups using two-dimensional GMM fitting (CA1-LECs vs. CA1-LECd). The optimal number of clusters was defined by the BIC (Figure S4C). These procedures were performed with custom scripts written in Python (ver. 3.9; Python Software Foundation, DE, US) along with some additional modules such as scikit-learn (Pedregosa et al., 2011; for 1D and 2D GMM fittings with EM algorithm and following BIC calculation) and PyClustering (Novikov, 2019; for *x*-means clustering).

### Histological observations

After the recording experiments, animals were deeply anesthetized with urethane (2–3 g/kg, i.p., Nacalai Tesque) and transcardially perfused with cold saline followed by 4% formaldehyde in 0.1 M phosphate buffer. Whole brains were post-fixed and sliced coronally into 50-μm serial sections using a microslicer (VT1000S, Leica, Wetzlar, Germany). Electrode tracks labeled with 1,1’-dioctadecyl-3,3,3’,3’-tetramethylindocarbocyanine perchlorate (DiI, Thermo Fisher Scientific, MA, USA) were observed in the CA1 and LEC under a fluorescence microscope (BX51N, Olympus).

### Retrograde tracing

Rats were anesthetized with isoflurane in an induction chamber and were moved to a surgical mask on a stereotactic frame. The skull was exposed and a small burr hole was drilled above the injection site. The injection was made by means of a glass micropipette (tip diameter = 20–40 µm) connected to a 1 µL Hamilton microsyringe. Rats received 100 nL of Fluoro-Gold (2.5% in H_2_O, Fluorochrome) and 1,200 nL of mRFP expressing G-deleted rabies viral vector (rHEP5.0-ΔG-mRFP; 6.0 × 10^8^ focus forming units (FFU)/mL; Ohara et al., 2013) into the mPFC (AP = +3.5; ML = 0.6; DV = −2.6) and the dorsal hippocampus (AP = −4.4; ML = 1.8; DV = −2.6), respectively. Injection site coordinates were based on the rat brain atlas (Paxinos and Watson, 2006) and calculated from bregma. Seven days into the survival period, rats were deeply anesthetized with sodium pentobarbital (100 mg/kg, i.p.) and perfused transcardially with Ringer’s solution (0.85% NaCl, 0.025% KCl, 0.02% NaHCO_3_) followed by 4% paraformaldehyde in 0.1 M phosphate buffer. Brains were removed from skulls, postfixed in 4% paraformaldehyde in 0.1 M phosphate buffer for 4 h at 4°C, and subsequently cryoprotected in a mixture of 20% glycerol and 2% dimethyl sulfoxide. The brains were cut into 40 µm sections in coronal plane on a freezing microtome. Sections were counterstained with mouse anti-NeuN antibody (Millipore #MAB377) as described previously (Ohara et al., 2021a), mounted on gelatin-coated slides, and covered with Entellan new (Millipore, #107961) before a coverslip was applied. Axio Scan. Z1 (Carl Zeiss) and ZEN 2 software (Carl Zeiss) were used to image labeled neurons.

### Experimental design and statistical analysis

We obtained electrophysiological data from 25 sessions in 25 Thy1-ChR2 rats to examine forelimb-movement representations of the CA1 and LEC neurons. In total, we included data from 829 CA1 neurons (pre-training, 296 cells; post-training, 533 cells), and 1,287 LEC neurons (pre-training, 370 cells; post-training, 917 cells) during task performance (≥20 trials with total ≥250 spikes; see Results for details). These neurons were divided into RS and FS subclasses by spike duration, and were further classified into Go-type, Go+Reward-type, Hold-type, Hold+Reward-type, and Reward-type neurons if they were related with task events functionally (Figures 3−7). Data in the text and figures are expressed as means ± SD (unless otherwise noted) and sample number (*N*). Sample sizes (the number of animals, sessions, and neurons) were estimated according to previous studies (Soma et al., 2017; 2019) and confirmed to be adequate by power analyses (power = 0.9; alpha error = 0.05). We used the following statistical methods: KS test, Mann–Whitney test, Wilcoxon signed-rank test, χ^2^ test with post hoc residual analysis, Kruskal-Wallis test with post hoc Steel-Dwass test, and two-way ANOVA. All tests were two-sided unless otherwise stated. These statistical tests were conducted with MATLAB’s Statistics and Machine Learning Toolbox (MathWorks). Differences were considered statistically significant when *p* < 0.05 (see Results for details). Blinding and randomization were not performed.

## Declaration of interests

The authors declare no competing interests.

## Author contributions

S.S., S.O., and Y.I. designed the research; S.S., S.O., S.N., J.Y., and E.P. performed the research; S.S., N.S., and Y.S. analyzed the data; and S.S., S.O., E.P., K.T., and Y.I. wrote the manuscript.

## Funding

This work was supported by Grants-in-Aid for Young Scientists (20K15934 to S.S.), for Scientific Research on Innovative Areas (20H05069 to S.S.; 16H01516 to Y.S. and Y.I.; 18H05524 to Y.S.; 20H05053 to Y.I.; 21H00178 to S.O.), for Scientific Research (19K06917 to S.O.), for Transformative Research Areas (A) (21H05242 to Y.I.) from JSPS; Brain/MINDS (JP19dm0207089 to Y.I.) from AMED; CREST (JPMJCR1751 to Y.I.), PRESTO (JPMJPR21S3 to S.O.) from JST; the Supported Program for the Strategic Research Foundation at Private Universities (S1311013 to Y.S. and Y.I.) from MEXT; the Shimizu Foundation for Immunology and Neuroscience Grant for 2019 (S.S.); and the Takeda Science Foundation (Y.I.).

## Acknowledgements

We are grateful to Drs. T. Samura, A Rios, M Kawabata, Y. Kawaguchi, M. Kimura, A. Nambu, and S. Okabe for helpful suggestions and discussion. We also thank M. Goto, C. Soai, and H. Yoshimatsu for technical assistance.

---

**Figure S1.**
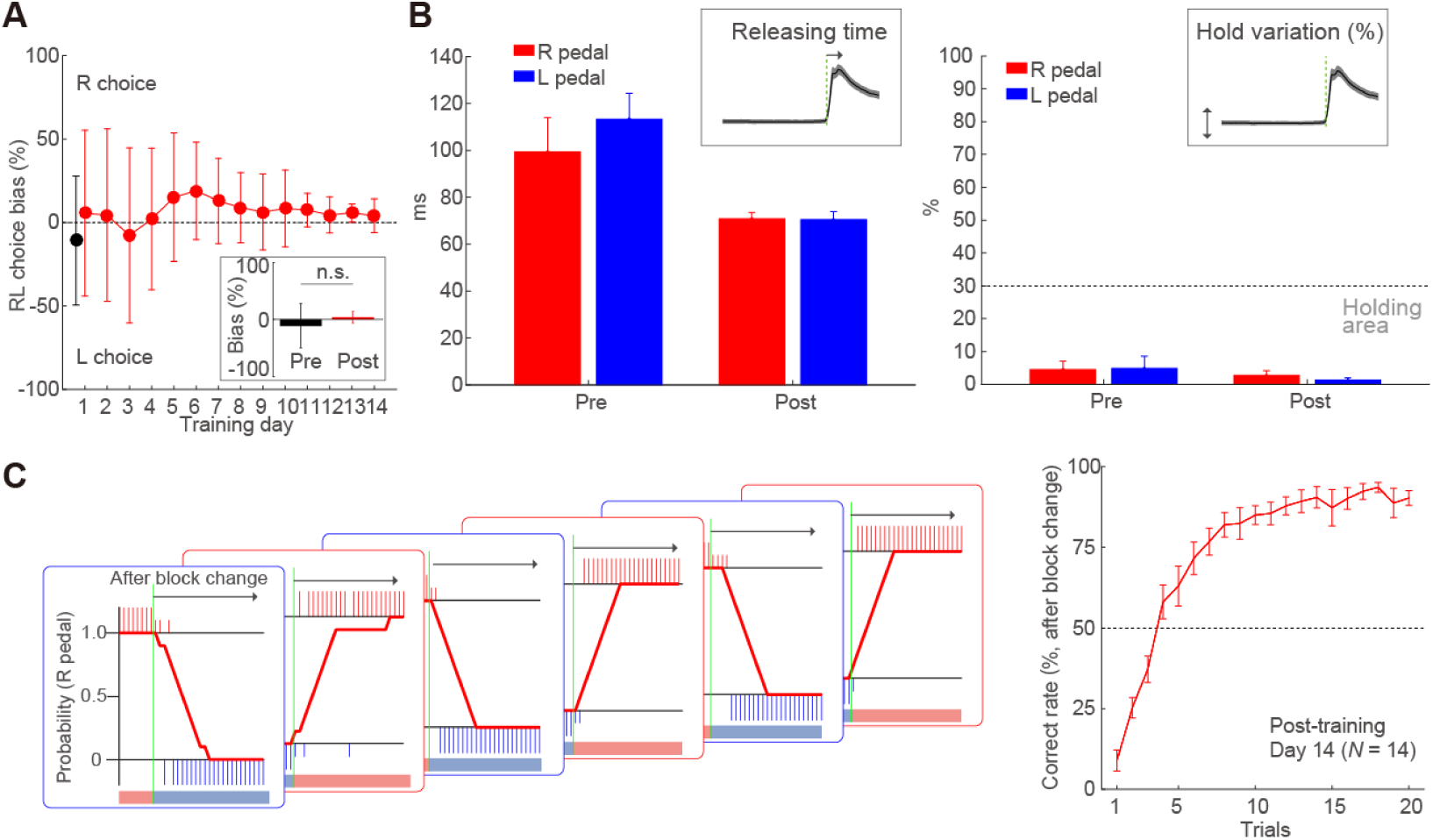
Behavioral task performance on the self-paced, spontaneous left or right pedal pressing task. (A) Left-right choice bias (RL choice bias). Both groups were able to manipulate the individual pedals spontaneously without left-right bias and there was no significant difference of bias of left-right pedal manipulation between groups (Inset, Mann– Whitney test, z = 1.23, p = 0.20, r = 0.26). (B) The stability of holding and releasing time (time from the onset to end of release) were developed over the training. The rats of the post-training group quickly released the pedal (releasing time [ms], group, F(1,49) = 17.9, *p* < 1.1 × 10^−4^, ηG^2^= 0.27; right vs left, F(1,49) = 0.50, *p* = 0.48, ηG^2^= 0.01) and returned their forelimb to the pedal after stable pedal holding (two-way ANOVA, group, F(1,49) = 16.1, *p* < 2.1 × 10^−4^, ηG2= 0.25; right vs left, F(1,49) = 0.97, *p* = 0.33, ηG^2^= 0.02). (C) Averaged proportion of correct choices in the first to 20^th^ trials after the change of blocks. The number of trials to achieve the reversal (> 50%) of the proportion of correct choices after the block change is 5.1 ± 1.9.

**Figure S2.**
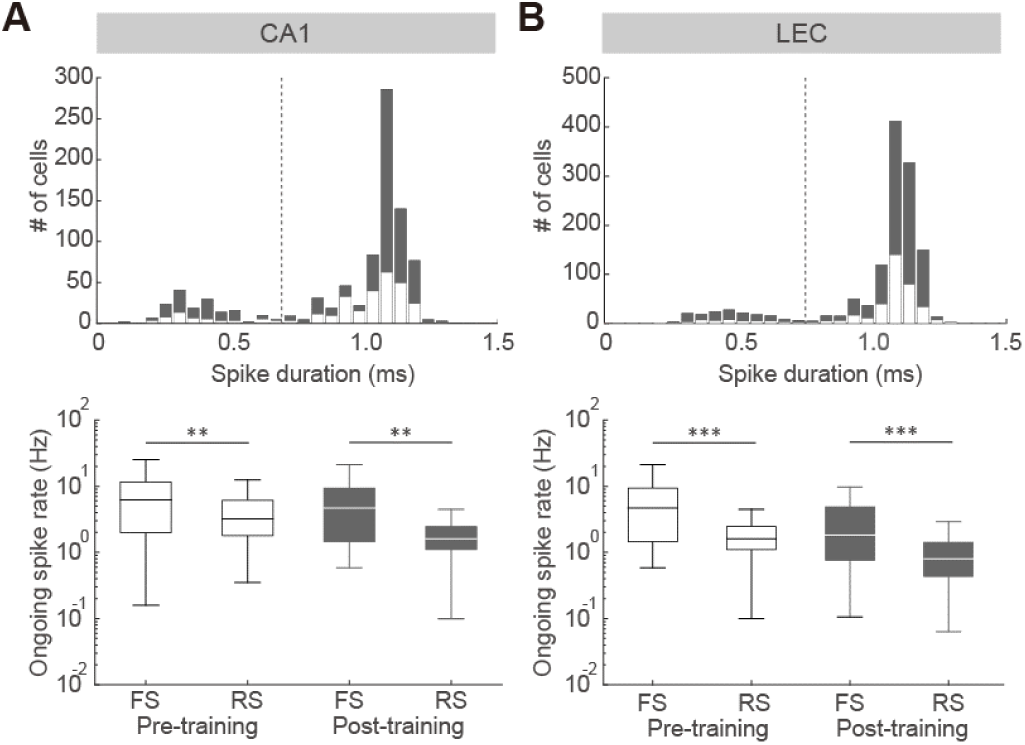
Classification of RS and FS neurons in the CA1 and LEC. (A and B) Recorded neurons were divided into regular-spiking (RS) and fast-spiking (FS) neurons based on spike duration (RS: ≥0.6 ms, triangles; FS: <0.6 ms, circles). The spike duration was defined as the time from spike onset to the first positive peak. Top, bimodal distribution of spike durations. White and gray colors represent neurons obtained from pre- and post-training groups, respectively. Bottom, box plot shows ongoing (all averaged) spike rate (middle bar, median; upper and lower edges, quartiles; whiskers, error) for FS and RS neurons in the CA1 (A) and LEC (B). The ongoing spike rates of FS subtypes were significantly higher than those of RS subtypes in the CA1 and LEC (Pre-training: CA1-RS, *n* = 247, mean ± SD, 4.4 ± 3.8 Hz, CA1-FS, *n* = 49, 7.8 ± 7.0 Hz; Mann– Whitney test, *z* = −2.97, *p* < 3.0 × 10^−3^, *r* = 0.17; LEC-RS, *n* = 335, 2.0 ± 1.4 Hz, LEC-FS, n = 35, 5.7 ± 4.9 Hz; *z* = −4.60, *p* < 4.2 × 10^−6^, *r* = 0.24; Post: CA1-RS, *n* = 433, 3.8 ± 4.0 Hz, CA1-FS, *n* = 100, 6.7 ± 7.9 Hz; *z* = −2.98, *p* < 2.9 × 10^−3^, *r* = 0.12; LEC-RS, *n* = 789, 1.2 ± 1.3 Hz, LEC-FS, *n* = 128, 4.4 ± 8.7 Hz; *z* = −7.38, *p* < 1.5 × 10^−13^, *r* = 0.24).** *p* < 0.01, *** *p* < 0.001, Mann–Whitney test.

**Figure S3.**
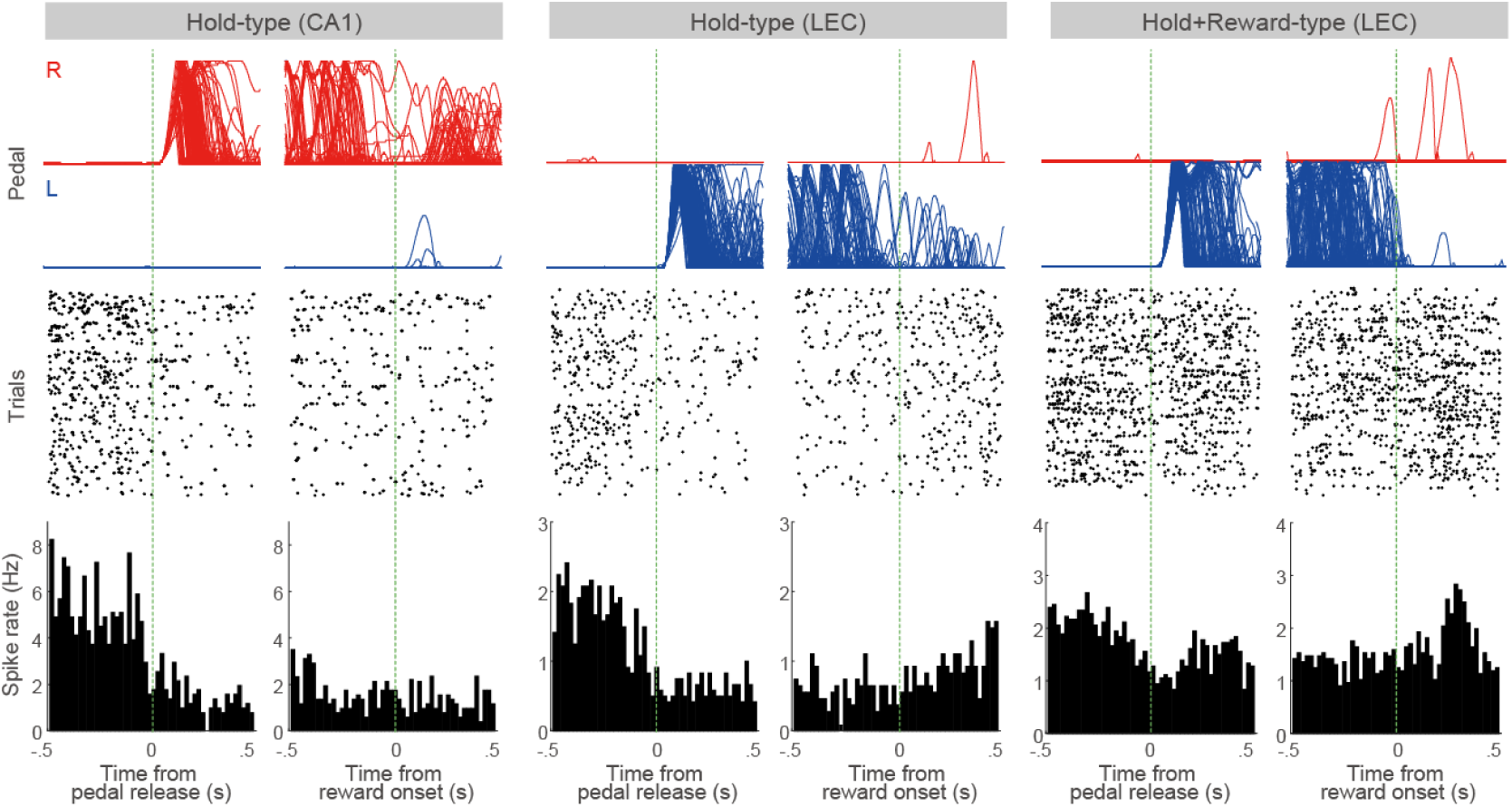
Example Hold-related activities of CA1 and LEC neurons during task performance. Hold-related activities after learning (left, Hold-type in the CA1; middle, Hold-type in the LEC; right, Hold+Reward-type in the LEC). Spikes were aligned with the onset of pedal release (left column) or reward delivery (right column) at 0 s for each cell. Top, pedal trajectories (R: right pedal, L: left pedal). Middle, raster plots. Bottom, peri-event time histograms (bin width, 20 ms).

**Figure S4.**
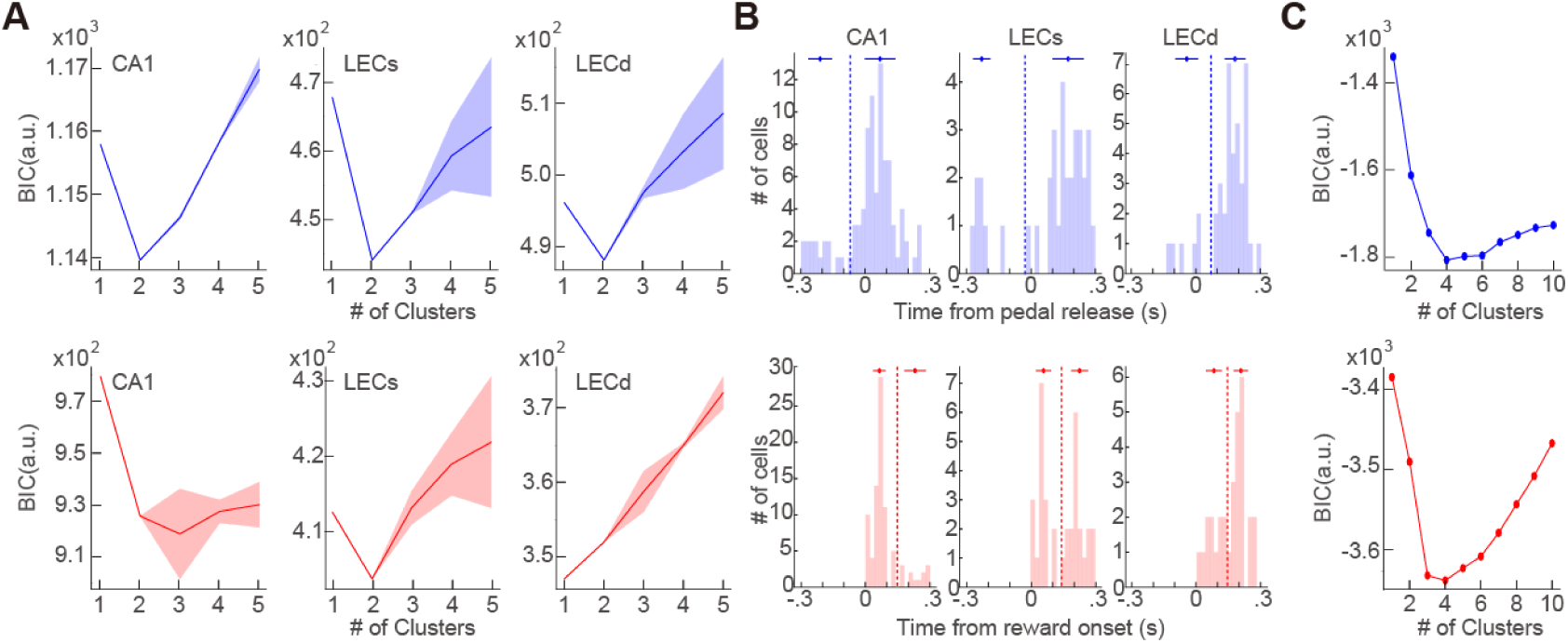
BIC values and Peak latency clustering with different methods. (A) BIC values for cluster numbers of 1 to 5, for the peak latency distributions (Figure 6C). Mean ± SD. Fitting with one or two clusters resulted in the least BIC. (B) *x*-means clustering of peak latencies in CA1 (left), and superficial (middle) and deep (right) layers of LEC, aligned with pedal release (top) and reward onset (bottom) timings. Symbols and horizontal lines indicate mean and ± SD of each cluster, respectively. Dotted vertical lines are boundaries of clusters. (C) BIC values for cluster numbers of 1 to 10, for the pseudo-paired latency difference distribution (Figure 6D). Fitting with four clusters resulted in the least BIC for both pedal-release and reward-onset alignment conditions.

**Figure S5.**
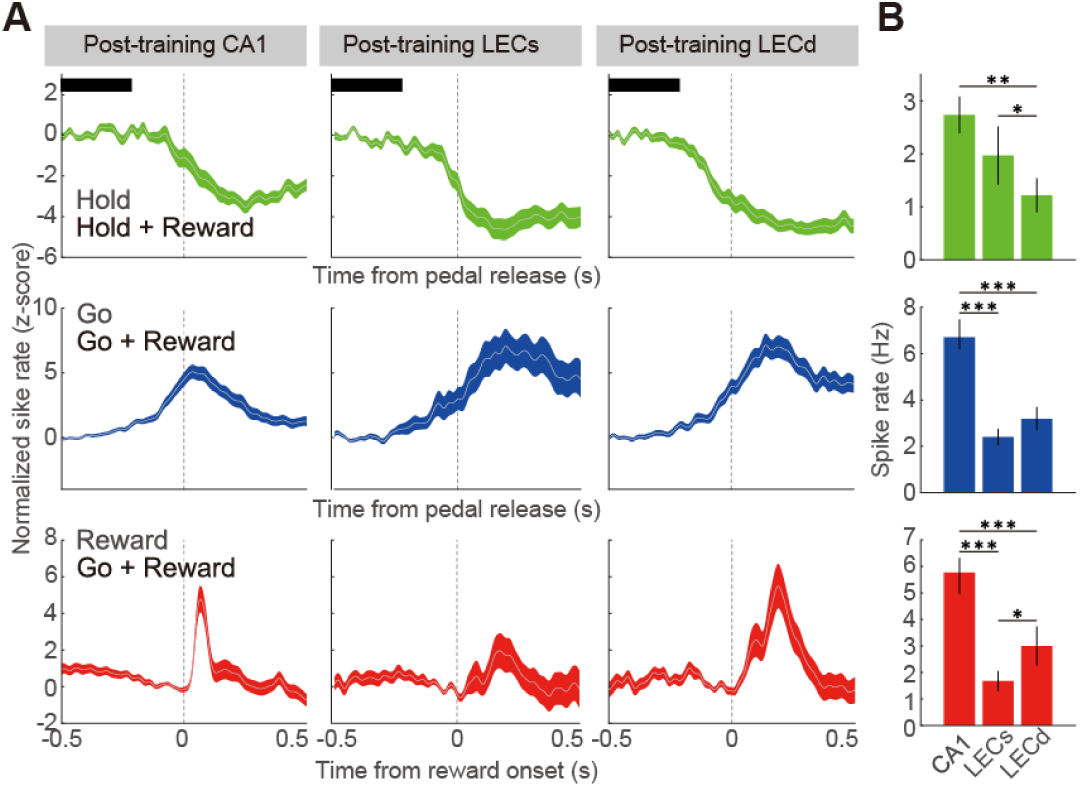
Differences of task-related activity among the CA1 and superficial and deep layers of the LEC. (A) Averaged PETHs of all hold-related (Hold-type and Hold+Reward-type), go-related (Go-type and Go+Reward-type), and reward-related (Reward-type and Go+Reward-type) activities in the CA1 (left), and superficial (middle) and deep layers of the LEC (right). PETHs were aligned with pedal release onset (top and middle) and reward onset at 0 s (bottom). Shaded regions represent 95% CIs. (B) Comparison of spiking activities among the CA1, and superficial and deep layers of the LEC. CA1 neurons showed higher activity than LEC neurons. Horizontal black bars in the Hold-related activity (A) indicate the windows for calculating the firing rate. The spiking rate for go- and reward-related activities were calculated from the peak period (peak ± 150 ms). In the hold-related activity, superficial layer neurons showed a higher spiking rate than deep layer neurons. In contrast, deep layer neurons showed a higher spiking rate than superficial neurons in the reward-related activity. * *p* < 0.05, ** *p* < 0.01, *** *p* < 0.001, post hoc Steel-Dwass test.

**Table S1:** Comparison of peak activity triggered by the pedal release between contra- and ipsilateral trials (Mann–Whitney test)

| Area | Activity type | Contra (Hz) | Ipsi (Hz) | z value | p value | Effect size (r) |
| --- | --- | --- | --- | --- | --- | --- |
| CA1 | Go-type | 6.2 ± 4.2 | 6.9 ± 4.6 | -3.07 | 2.1 × 10<sup>-3</sup> | 0.30 |
|  | Go+Reward-type | 9.6 ± 6.8 | 9.9 ± 7.2 | -0.83 | 0.40 | 0.10 |
| LECs | Go-type | 2.2 ± 2.0 | 2.0 ± 1.3 | 0.70 | 0.48 | 0.10 |
|  | Go+Reward-type | 3.0 ± 2.6 | 2.8 ± 2.4 | -1.10 | 0.30 | 0.10 |
| LECd | Go-type | 3.1 ± 2.4 | 2.7 ± 2.4 | 1.53 | 0.13 | 0.20 |
|  | Go+Reward-type | 4.3 ± 3.7 | 5.2 ± 3.4 | -0.73 | 0.50 | 0.10 |

Values are means ± SD

**Table S2:** Comparison of peak activity triggered by the reward onset between contra- and ipsilateral trials (Mann–Whitney test)

| Area | Activity type | Contra (Hz) | Ipsi (Hz) | z value | p value | Effect size (r) |
| --- | --- | --- | --- | --- | --- | --- |
| CA1 | Go-type | 4.6 ± 3.6 | 4.8 ± 3.5 | -0.70 | 0.48 | 0.10 |
|  | Go+Reward-type | 9.4 ± 7.3 | 9.2 ± 7.0 | -1.73 | 0.08 | 0.18 |
| LECs | Go-type | 1.3 ± 0.9 | 1.5 ± 1.3 | -1.49 | 0.15 | 0.10 |
|  | Go+Reward-type | 2.9 ± 2.6 | 2.6 ± 2.5 | -0.51 | 0.61 | 0.08 |
| LECd | Go-type | 1.3 ± 1.0 | 1.6 ± 1.0 | -0.87 | 0.41 | 0.08 |
|  | Go+Reward-type | 4.2 ± 3.7 | 5.1 ± 3.6 | -1.16 | 0.25 | 0.20 |

Values are means ± SD

## Notes

### Competing Interest Statement

The authors have declared no competing interest.

